# Pneumolysin-dependent and independent non-canonical autophagy processes mediate host defense against pneumococcal infection

**DOI:** 10.1101/2025.02.11.637156

**Authors:** Bartosz J. Michno, Niedharsan Pooranachandran, Tonisha C. Smith, Erin Faught, Sandra Lipowská, Andrew K. Fenton, Annemarie H. Meijer, Tomasz K. Prajsnar

**Affiliations:** Department of Evolutionary Immunology, Institute of Zoology and Biomedical Research, Faculty of Biology, Jagiellonian University, Krakow, Poland; Doctoral School of Exact and Natural Sciences, Jagiellonian University, Krakow, Poland; School for Biosciences, Florey Institute for Host-Pathogen Interactions, University of Sheffield, Sheffield, United Kingdom; Institute of Biology Leiden, Leiden University, Leiden, The Netherlands

**Keywords:** CASM, LAP, macrophage, *pneumoniae*, STIL, *Streptococcus*, zebrafish

## Abstract

*Streptococcus pneumoniae* is an opportunistic pathogen responsible for life-threatening diseases including pneumonia and meningitis. The host defense against pneumococci relies heavily on macrophages, which can effectively internalize and degrade bacteria. Recent studies have implicated both canonical and non-canonical autophagy-related processes in bacterial clearance, but the precise pathways mediating defense against *S. pneumoniae* remain unknown. Here, we utilize a well-established zebrafish larval infection model to investigate the role of autophagy in host defense against pneumococci *in vivo*. Using a transgenic macroautophagy/autophagy reporter line, we found the autophagy marker Map1lc3/Lc3 being recruited to pneumococci-containing vesicles upon bacterial internalization by zebrafish macrophages. The genetic inhibition of core autophagy gene *atg5* led to loss of the Lc3 associations and their impaired acidification, significantly delaying bacterial clearance. This Lc3 recruitment is partially mediated by LC3-associated phagocytosis (LAP), as knockdown of *cyba* and *rubcn* moderately reduced Lc3 association with phagosomes and diminished pneumococcal degradation. Interestingly, we observed no involvement of xenophagy components in *S. pneumoniae*-infected macrophages, suggesting the activation of another non-canonical autophagy pathway, distinct from LAP, targeting pneumococci-containing phagosomes. Instead, we found that the pneumococcal pore-forming toxin pneumolysin induces ROS-independent CASM pathways, one of which is abolished by knockdown of *tecpr1a* indicating the involvement of sphingomyelin-Tecpr1-induced LC3 lipidation (STIL). Collectively, our observations shed new light on the host immune response against *S. pneumoniae*, demonstrating that several distinct non-canonical autophagy pathways mediate bacterial degradation by macrophages and providing potential targets for the development of novel therapies to combat pneumococcal infections.

## Introduction

*Streptococcus pneumoniae* (pneumococcus) is an opportunistic pathogen that colonizes the human upper respiratory tract and is able to invade other tissues including the central nervous system. This bacterium causes several life-threatening diseases such as community-acquired pneumonia [1] resulting in more than 1 million deaths worldwide each year [2]. Pneumococcus has been repeatedly included in the WHO’s list of priority bacterial pathogens since 2017 to date (WHO report, 2024). The widespread use of various antibacterial drugs, mostly β-lactam antibiotics such as penicillin or ampicillin, combined with the high genome plasticity of pneumococci facilitates the spread of pneumococcal resistance to drugs and vaccines, limiting their effectiveness [3,4]. Therefore, alternative treatment strategies are urgently needed to combat this pathogen.

The first line of defense against invading pneumococci are tissue-resident alveolar macrophages in the lung [5]. Once internalized, the bacteria sequestered in phagosomes are rapidly acidified by activation of the vacuolar-type H^+^-translocating ATPase (V-ATPase), which pumps protons into the vacuole [6–8]. This acidification is required for *S. pneumoniae* killing by phagocytes in the zebrafish model system [9]. Furthermore, lysosomal-dependent degradation, following fusion of phagosomes with lysosomes, has been well described in the literature and is known to be involved in the antibacterial response [10]. Aside from classical phagocytosis, other pathways can also lead to intracellular bacteria degradation including autophagy-related mechanisms which are currently being intensively studied [11]. These are either canonical pathogen-selective autophagy, also known as xenophagy [12], or more recently identified non-canonical autophagy pathways, collectively referred to as conjugation of ATG8 to single membranes (CASM) [13,14].

Xenophagy is a form of canonical autophagy in which either ruptured bacteria-containing phagosomes or pathogens that have escaped into the cytosol are detected and tagged by various proteins such as ubiquitin or glycan-binding galectins [15,16]. Subsequently, such labeled bacteria bind to sequestosome-like receptors (SLRs) and MAP1LC3/LC3 (microtubule-associated protein 1 light chain 3) [12,17,18]. LC3 association promotes the phagophore expansion resulting in formation of a double-membrane autophagosome [19]. It has recently been reported that SLRs such as SQSTM1/p62 (sequestosome 1) and CALCOCO2/NDP52 (calcium binding and coiled-coil domain 2) contribute to the initiation of xenophagy of intracellular *S. pneumoniae* within non-professional phagocytes such as human pulmonary epithelial cells and mouse embryonic fibroblasts (MEFs) *in vitro* [20,21]. It has been also shown that pneumolysin (Ply), a pore-forming toxin produced *by S. pneumoniae* that is able to induce membrane rupture of the vesicles containing bacteria leading to subsequent xenophagy [21].

The most studied non-canonical autophagy pathway is LAP, a form of CASM that involves many of the same molecular components as xenophagy, but not SLRs. In this case, LC3 is recruited to the single-membrane phagosome triggered by perturbation of ionic and pH balance [13]. It is generally accepted that LC3 lipidation in LAP requires the activity of NOX (NADPH oxidase) and CYBA, generating reactive oxygen species (ROS) [22]. This ROS production has been shown to promote an interaction between the V-ATPase and ATG16L1, which initiates LC3 lipidation [23]. We have recently shown that once ROS production is blocked, the LAP response to various bacterial pathogens is greatly reduced [24,25]. In addition, RUBCN/RUBICON (rubicon autophagy regulator) has been implicated as a positive regulator of LAP acting upstream of NADPH oxidase complex assembly [22]. Recent *in vitro* evidence has shown that ROS-dependent LAP occurs within mouse bone marrow-derived macrophages (BMDMs) infected by pneumococci and significantly contributes to bacterial degradation [26].

Although the degradation of invading *S. pneumoniae* through LAP or xenophagy has been investigated *in vitro*, results differ between studies in professional and non-professional phagocytes. While in MEFs, xenophagy has been demonstrated to play a role in pneumococcal degradation and its activation was preceded by atg16l1-dependent but ROS-independent LAP-like LC3 associations [20,21], no such response was observed in macrophages [26]. Additionally, the specific interplay of these pathways in the wider and more complex host environment have not yet been elucidated. Unveiling the role of these mechanisms *in vivo* would support the development of innovative immunomodulatory approaches against *S. pneumoniae* in the future.

Previously, we established a model of pneumococcal infection in larval zebrafish and described that zebrafish macrophages can prevent dissemination of intravenously administered unencapsulated *S. pneumoniae* [9]. The macrophages protect infected larvae by internalization and acidification-mediated killing of pneumococci, underscoring the cross-species evolutionary conserved role of these professional phagocytes in host protection against this pathogen, and providing new opportunities to gain insight into the host-pneumococcus interactions *in vivo*.

In the present study, we focus on the role of autophagy-related mechanisms in *S. pneumoniae* infection using zebrafish larvae. We demonstrate that upon internalization of pneumococci, the vast majority of infected macrophages rapidly respond by the Lc3 conjugation to phagosomal membrane which facilitates phagosomal acidification and bacterial killing. We also show that, rather than being mediated by xenophagy, the protective role of autophagy in pneumococcal infection of macrophages is primarily associated with CASM such as LAP and other pneumolysin-mediated pathways including the sphingomyelin-TECPR1-induced LC3 lipidation (STIL).

## Results

### Intracellular *S. pneumoniae* triggers Lc3 recruitment to bacteria-containing phagosomes in zebrafish macrophages

To determine whether the autophagy response occurs to intracellular pneumococci, we used our previously established *S. pneumoniae* infection model of zebrafish larvae [9] using the transgenic zebrafish line *Tg(CMV:EGFP-map1lc3b)^zf155^*, hereafter called *CMV:GFP-Lc3* [27]. The GFP-LC3 reporter has been widely used in studying various autophagy pathways in response to infection [28,29]. The zebrafish larvae were injected intravenously with fluorescently labeled unencapsulated *S. pneumoniae* (D39 Δ*cps hlpA*::*mKate2*) and the infected larvae fixed at 2, 4 and 7 h post infection (hpi). Confocal analysis revealed robust activation of the autophagy machinery, as more than 70% of infected phagocytes contained bacterial clusters surrounded by Lc3 at 2 hpi and this response gradually decreased by 7 hpi (Fig. 1A-B). The time-dependent loss of Lc3-bacteria associations is probably associated with rapid autophagic flux leading to degradation of internalized pneumococci upon phagosomal acidification that we have observed before [9]. Indeed, in addition to the decrease of Lc3 associations over time, the distinctive coccus shape of internalized fluorescent bacteria was also lost as a result of subsequent degradation (Fig. 1A). To determine whether this autophagic response is not specific to the capsule-negative D39 Δ*cps* strain, we also analyzed the infection with the parental encapsulated wild-type (D39 WT) *S. pneumoniae* and observed comparable levels of the autophagic machinery activation despite less effective phagocytosis of D39 WT (Fig. S1A-B). Interestingly, the observed high levels of the Lc3-association response to *S. pneumoniae* infection within zebrafish macrophages contrasts with the response to other Gram-positive bacterium - *Staphylococcus aureus* [25], where the Lc3 response occurs less frequently.

**Figure 1.**
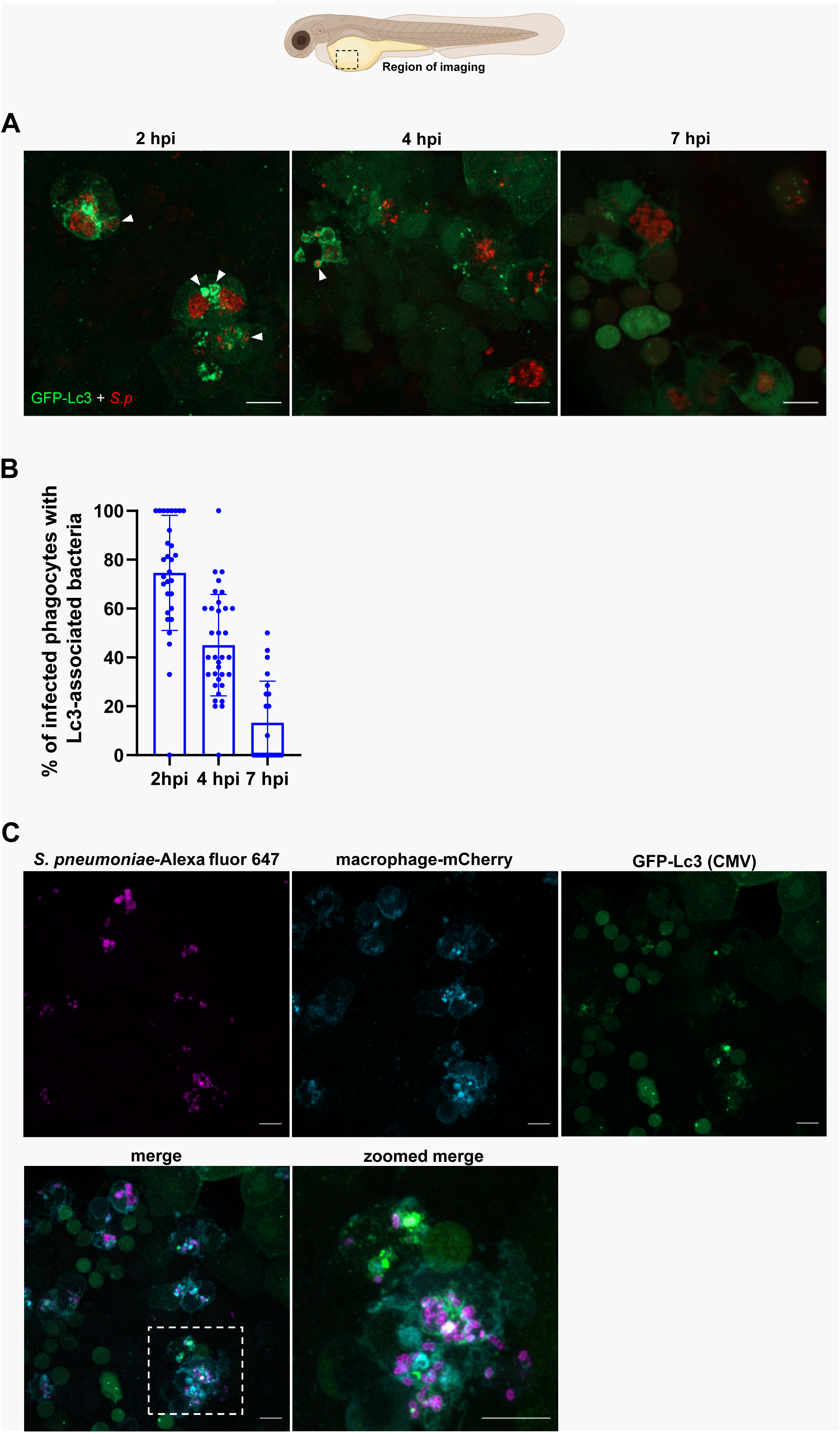
The autophagic response occurs during systemic infection with pneumococci. (**A**) representative confocal images of *CMV:GFP-Lc3* larvae infected systemically with approximately 1600 CFU of mKate2-labeled D39 Δ*cps S. pneumoniae* and fixed at 2, 4 and 7 hpi. Scale bars: 10 µm. Arrowheads indicate Lc3-positive vacuoles containing bacteria. (**B**) Quantification of Lc3 associations with intracellular *S. pneumoniae* within infected phagocytes of fixed *CMV:GFP-Lc3* at 2, 4 and 7 hpi. Data are shown as mean ± standard deviation (SD). Data obtained from two independent experiments (**C**) representative confocal images of *CMV:GFP-Lc3 mpeg:mCherry-F* dual transgenic larvae infected systemically with approximately 1600 CFU of Alexa Fluor 647-stained D39 Δ*cps S. pneumoniae* and fixed at 2 hpi.

Since we have previously shown that the vast majority of unencapsulated pneumococci are internalized by macrophages [9], we made an assumption that the observed Lc3-positive infected phagocytes are indeed macrophages. Nevertheless, to confirm the identity of these cells we crossed the *CMV:GFP-Lc3* zebrafish with the *Tg(mpeg1:mCherryF)^ump2^* transgenic line (hereafter called *mpeg:mCherry-F*), where the membrane-localized mCherry reporter is expressed in zebrafish macrophages [30]. The resulting dual transgenic larvae were intravenously infected with Alexa Fluor 647-labelled pneumococci. Microscopy analysis confirmed that out of 63 observed phagocytes evoking the Lc3-mediated response to pneumococci, 62 were actually macrophages (98.4%) (Fig. 1C). Altogether, these results demonstrate that larval zebrafish serve as a suitable *in vivo* model to study intramacrophage autophagic response to pneumococci.

### Genetic inhibition of autophagy impairs pneumococcal killing

Having identified the Lc3 response to *S. pneumoniae*, we subsequently asked whether this process is actively involved in bacterial clearance or helps infected phagocytes to eliminate invading pneumococci by some alternative mechanism.

To this end, using CRISPR-Cas9 we knocked down the autophagy-essential gene *atg5,* encoding protein that facilitates the conjugation of phosphatidylethanolamine to the activated Lc3 [31]. The knockdown efficacy was verified by PCR and restriction fragment length polymorphism/RFLP analysis (Fig. S2A-B). We found that the Lc3 signal surrounding the internalized bacteria was greatly reduced (Fig. 2A-B), which indicates that the observed Lc3 associations are indeed the result of an autophagy response to internalized pathogens. Next, the infected control and *atg5* knockdown larvae were homogenized and the number of bacteria within individual larvae were enumerated over the course of the infection. We found that although vast majority of bacteria were cleared by 7 hpi in both groups, and all infected control and *atg5* larvae survived until 120 hpf (Fig. S9A), the larvae with impaired autophagy machinery required more time to eliminate the pathogens, especially in the early stages of infection (Fig. 2C). The observation that *S. pneumoniae* clearance is dampened within zebrafish with an impaired autophagic response suggests a host-protective role for the process of Lc3-bacteria association. Previously, we have shown that acidification is required for *S. pneumoniae* killing by phagocytes in the zebrafish model system (Prajsnar et al., 2022). Therefore, in order to determine whether the observed autophagic response plays a role in phagosomal acidification, we injected pneumococci pre-stained with a pH-sensitive dye - pHrodo Red into control and autophagy-deficient zebrafish larvae. We observed significantly elevated levels of bacteria within acidic compartments of zebrafish macrophages of control larvae in comparison to the *atg5-* and *atg16l1*-depleted counterparts (Fig. 2D-E). Together, these data show that the autophagic response to internalized *S. pneumoniae* enhances the pace of pneumococcal killing by increasing the acidification rate of the bacteria-containing vesicles.

**Figure 2.**
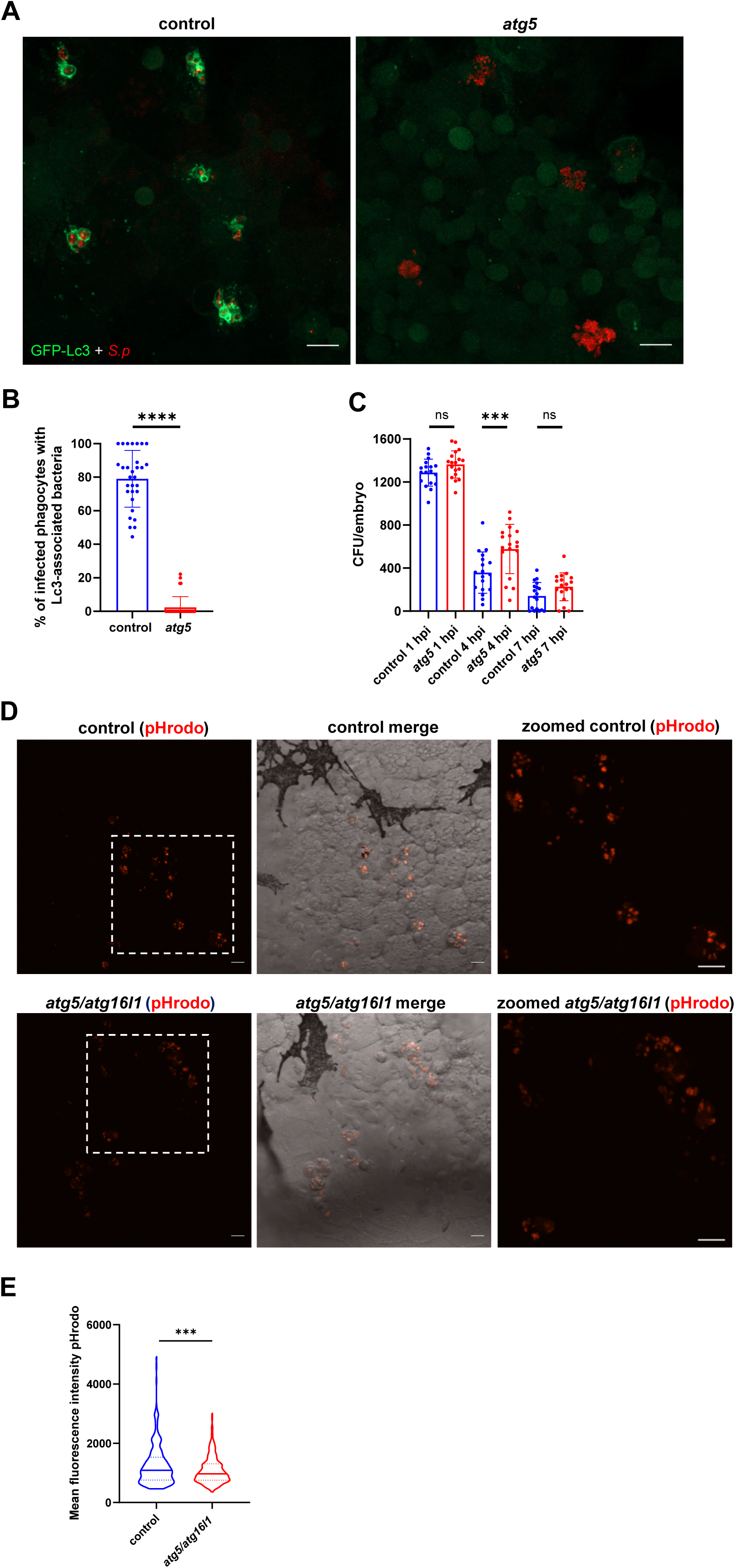
The autophagic response contributes to phagosomal acidification and pneumococcal clearance. (**A**) representative confocal images of control (left panel) and *atg5* knockdown (right panel) *CMV:GFP-Lc3* larvae infected systemically with approximately 1600 CFU of mKate2-labeled D39 Δ*cps S. pneumoniae* and fixed at 2 hpi. Scale bars: 10 µm. (**B**) Quantification of Lc3 associations with intracellular bacteria of control and *atg5* knockdown larvae. Data are shown as individual values ± standard deviation (SD). Data obtained from two independent experiments. (**C**) Numbers of viable colony forming units (CFUs) control or *atg5* knockdown larvae infected intravenously with approximately 1600 CFU of *S. pneumoniae* monitored at 1, 4, and 7 hpi. Data obtained from two independent experiments (**D**) Representative *in vivo* images of circulation valley of control (top) and *atg5*, *atg16l1* knockdown larvae (bottom) at 2 hpi injected with *S. pneumoniae* D39 Δ*cps* prestained with pHrodo Red. Panels on the right represent zoomed-in areas (indicated by dashed squares). Scale bars represent 10 µm . (**E**) Quantification of pHrodo Red fluorescence intensity within phagocytes containing pneumococci. Data obtained from two independent experiments). **** - p<0.0001, *** - p<0.001, ns - not significant

### Inhibition of reactive oxygen species (ROS) results in partial reduction of Lc3 associations and impaired bacterial killing

As we observed high levels of Lc3-positive vesicles within pneumococcus-infected macrophages, we sought to elucidate the molecular mechanism responsible for their formation. It is generally accepted that LC3 association with internalized pathogens can be mediated by either xenophagy or non-canonical autophagic pathways such as LAP. To determine which pathway is primarily responsible in our model, we manipulated phagosomal ROS production, as its inhibition prevents the LAP response in zebrafish to several pathogens [24,25], and recent findings suggest that ROS-dependent LAP is primarily responsible for pneumococcal clearance in BMDMs *in vitro* [26]. Using a previously validated morpholino-modified antisense oligonucleotide method [32], we inhibited the expression of *cyba/p22phox* (cytochrome b-245 alpha chain), a membrane-bound subunit of phagocyte NOX. While the morpholino-based targeting of *cyba* was previously shown to effectively inhibit Lc3 assembly on phagosome membranes during infection caused by either Gram-positive or negative bacteria [24,25], Fig. S1C-D), we surprisingly found that loss of *cyba* resulted in only partial reduction of Lc3 associations (Fig. 3A-B). The same Lc3 association pattern was observed in response to capsule-positive (D39 WT) pneumococci suggesting no role of pneumococcal capsule in autophagy induction (Fig. S1A-B).

**Figure 3.**
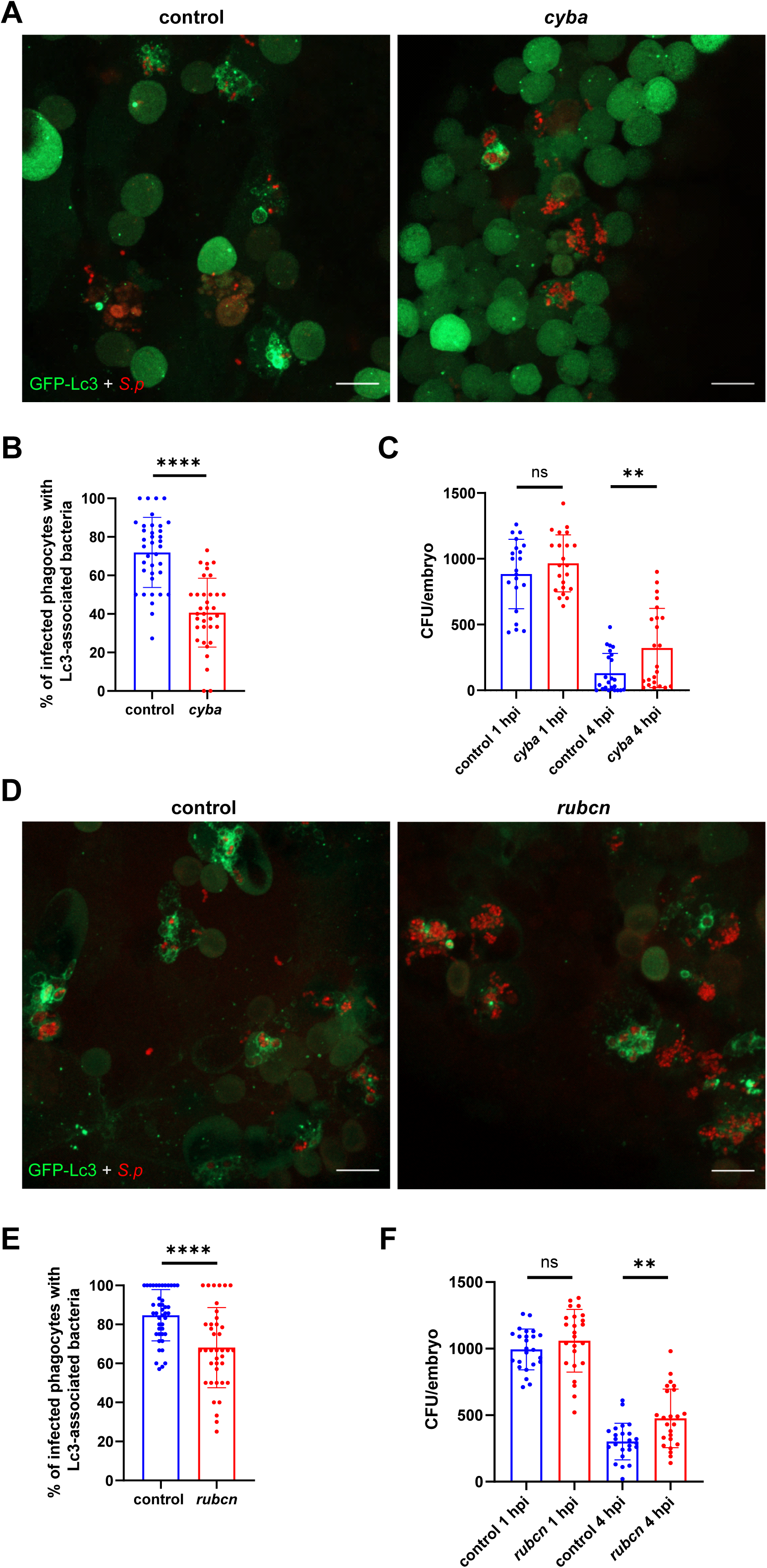
ROS-dependent autophagic response contribute to clearance of *S. pneumoniae*. (**A, D**) representative confocal images of control and indicated gene knockdown (A – *cyba*, D - *rubcn*) *CMV:GFP-Lc3* larvae infected systemically with approximately 1600 CFU of mKate2-labeled D39 Δ*cps S. pneumoniae* and fixed at 2 hpi. Scale bars: 10 µm. (**B, E**) Quantification of Lc3 associations with intracellular *S. pneumoniae* within infected phagocytes of fixed *CMV:GFP-Lc3* control and indicated gene knockdown (B – *cyba*, E - *rubcn*) larvae at 2 hpi. Data are shown as mean ± standard deviation (SD). Data obtained from three independent experiments. (**C, F**) Numbers of viable colony forming units (CFUs) over time of control and indicated gene knockdown (C – *cyba*, F - *rubcn*). Data obtained from three independent experiments.

In agreement with this genetic inactivation, chemical inhibition of NOX activity with diphenyleneiodonium (DPI) also showed a similar pattern of incomplete suppression of Lc3 aggregation with internalized pneumococci (Fig. S3). Another protein responsible for LAP is Rubicon, which is part of the class III phosphatidylinositol 3-kinase/PtdIns3K complex but also activates NOX promoting LAP [22,33]. Therefore, we knocked down *rubcn* (the knockdown efficacy was validated by qPCR followed by melt curve analysis for each targeted locus) and observed a significant decrease in Lc3 associations, but again, a large fraction of engulfed pneumococci were sequestered in LC3-positive vesicles (Fig. 3D-E) suggesting the existence of another autophagic pathway. To assess the function of LAP in bacterial killing, we enumerated the bacterial load in control and LAP-deficient infected larvae concentrating at early stages of infection as these timepoints showed significant differences when global autophagy was inhibited (Fig. 2C). Both *cyba* and *rubcn* knockdowns had significantly higher colony-forming units (CFU) counts at 4 hpi than in control siblings (Fig 3C,F). Taken together, these findings suggest that ROS- and Rubcn-dependent LAP occurs in response to pneumococci, plays a host-protective role during infection, but contributes only to a part of the autophagic response to pneumococci in zebrafish macrophages.

### Heat-inactivated or pneumolysin-deficient *S. pneumoniae* induce a diminished autophagic response within macrophages

Having established that LAP occurs in response to *S. pneumoniae* infection, we next tested whether the internalized bacterial cells need to be alive, and possibly secreting virulence factors, to induce the observed autophagic response.

First, to determine whether live *S. pneumoniae* cells are required to induce an Lc3-mediated response, larvae were injected with heat-killed bacterial cells. We observed a significant decrease in Lc3 associations with pre-killed bacteria (Fig. 4A-B). To further investigate whether LAP is involved in the response to heat-inactivated pneumococci, we injected *cyba* morphants with heat-killed *S. pneumoniae*. This combination led to an even more pronounced decrease in Lc3 recruitment to engulfed bacteria (Fig. 4A-B) indicating that LAP is evoked regardless of pneumococcal viability. Having shown that live pneumococci are required to fully induce the autophagic response in zebrafish macrophages, we speculated whether specific extracellular virulence factors play a role in this process. It has been demonstrated that the induction of the autophagic response to internalized pneumococci by professional and non-professional phagocytes *in vitro* requires Ply (Inomata et al., 2020; Ogawa et al., 2018, 2020).To test whether Ply is responsible for triggering autophagy in zebrafish macrophages, we first generated a *S. pneumoniae* strain in which the ply gene has been removed (Δ*ply*) and verified the absence of Ply protein expression by immunoblotting (Fig. 4C). We observed significantly less Lc3 associations in phagocytes when *S. pneumoniae* cells lack Ply compared to the control strain (Fig. 4D-E). It is worth noting that both heat-killed and Ply-deficient bacteria were still able to evoke a partial autophagic response (Fig. 4), similar to what we observed in ROS-depleted larvae (Fig. 3). These observations prompted us to perform an experiment where *cyba* morphants were injected with Ply-negative pneumococci. Strikingly, this combination led to a near-complete loss of Lc3 recruitment to internalized pneumococci (Fig. 4F-G), indicating that pneumolysin induces another autophagic pathway. Importantly, Ply-deficient pneumococci were phagocytosed by macrophages as effective as the parental strain (Fig. S9A-B), indicating that diminished autophagic response was not due to fewer intracellular bacteria. Interestingly, the bacteria without pneumolysin were also eliminated significantly faster by infected larvae than Ply-positive *S. pneumoniae* (Fig. S9C) as previously shown in a mouse model [34]. Additionally, we determined the role of pneumolysin in evoking inflammatory response in infected phagocytes. Zebrafish larvae of *il1b:GFP* [35] and *tnf*α*:GFP* [36] transgenic reporter lines infected with pneumolysin*-*deficient bacteria displayed higher expression of both pro-inflammatory cytokines in comparison to the parental strain (Fig. S9D-G). This stays in line with a recent report describing that pneumolysin can suppress the macrophage pro-inflammatory activation, contributes to immune evasion and bacterial persistence by modulating host immune responses [34].

**Figure 4.**
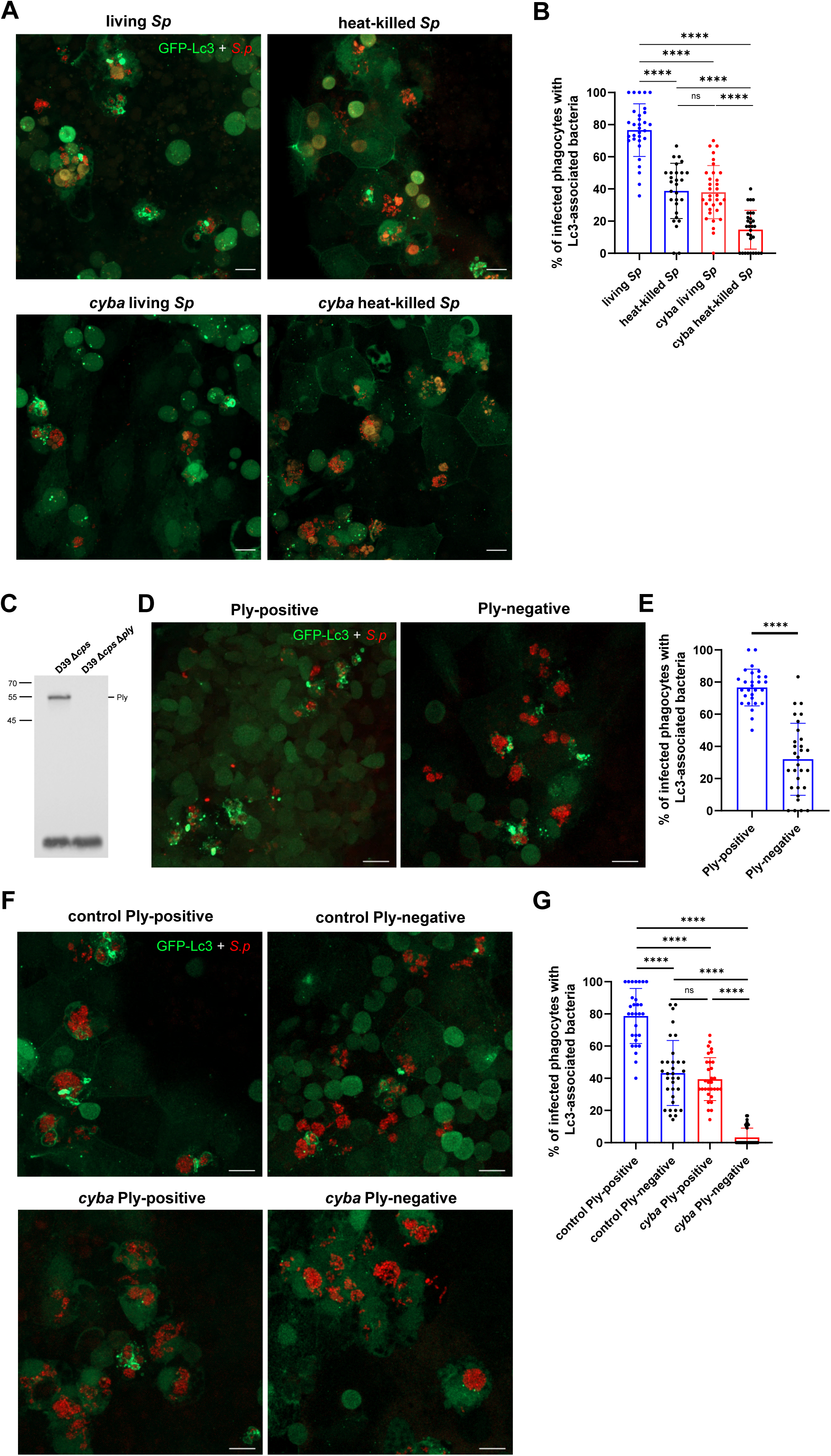
Live and pnemolysin-positive pneumococci are required to fully induce Lc3-mediated response within zebrafish macrophages. (**A**) representative confocal images of *CMV:GFP-Lc3* control larvae (top two panels) or *cyba* knockdown (bottom two panels) infected systemically with approximately 1600 CFU of live (left two panels) and heat-killed (right two panels) mKate2-labeled D39 Δ*cps S. pneumoniae* fixed at 2 hpi. Scale bars: 10 µm. (**B**) Quantification of Lc3 associations with intracellular live or heat-killed *S. pneumoniae* within infected phagocytes of fixed *CMV:GFP-Lc3* control or *cyba* larvae at 2 hpi. Data are shown as mean ± standard deviation (SD). Data obtained from two independent experiments. (**C**) Anti-Ply immunoblot analysis of whole-cell lysates from the indicated *S. pneumoniae* strains. The position of protein markers are indicated in kiloDaltons, Pneumolysin (Ply) ≈ 53.7 KDa. Representative blot shown, n = 4. (**D**) representative confocal images of control *CMV:GFP-Lc3* larvae infected systemically with approximately 1600 CFU of Ply-positive (D39 Δ*cps*, left panel) and Ply-negative (D39 Δ*cps* Δ*ply*, right panel) mKate2-labeled *S. pneumoniae* fixed at 2 hpi. Scale bars: 10 µm. (**E**) Quantification of Lc3 associations with intracellular Ply-positive or Ply-negative *S. pneumoniae* within infected phagocytes of fixed *CMV:GFP-Lc3* larvae at 2 hpi. Data are shown as mean ± standard deviation (SD). Data obtained from two independent experiments.

Overall, our observations confirm the role of pneumolysin in evoking an autophagic response in an in vivo system and importantly this pneumolysin-evoked response is independent of LAP.

### Lack of xenophagy components does not affect the autophagic response to pneumococci

We considered that xenophagy might be the Ply-mediated autophagy pathway, which complements the ROS-dependent LAP response. During xenophagy, ruptured phagosomes or cytosolic bacteria are detected and labeled with ubiquitin [20,37–39]. Furthermore, the expression of a pore forming toxin, ESAT-6, is required for inducing xenophagy in infection with mycobacteria [40]. To elucidate whether xenophagy is responsible for the pneumolysin-evoked and LAP-independent Lc3 response, we first sought to detect ubiquitinated bacteria. However, when we performed immunostaining for ubiquitin at 2 hpi, we observed ubiquitin staining in only around 5% of phagocytes containing Lc3-decorated bacteria, suggesting little involvement of pneumococcal ubiquitination within zebrafish phagocytes (Fig. S4).

Next, we took advantage of a previously validated double *sqstm1*^-/-^ *optn*^-/-^ mutant zebrafish line [39,41], lacking two key SLRs, Sqstm1/p62 (sequestosome 1) and optineurin (Otpn), whose contribution to xenophagy is firmly established and evolutionary conserve [42]. Using the CMV:GFP-Lc3 transgenic background, our confocal analysis revealed a comparable number of Lc3-positive infected phagocyted in the double mutant and wild-type larvae (Fig. 5A-B). Consistently, the kinetics of bacterial clearance remained virtually unchanged in Sqstm1- and Optn-deficient larvae when compared to the wild-type siblings (Fig. 5C). We also infected *sqstm1^-/-^* and *optn^-/-^* single mutants, and as expected, no differences in Lc3 formation or pneumococcal killing ability was observed (Fig. S5). Next, we reasoned that xenophagy might be triggered within *S. pneumoniae*-infected macrophages under conditions when ROS-dependent LAP is inhibited. To address this possibility, we knocked down *cyba* in *sqstm1*^-/-^ larvae, but found a similar reduction in Lc3 associations within infected phagocytes as those caused by ROS inhibition alone, without any further reductions (Fig. S6).

**Figure 5.**
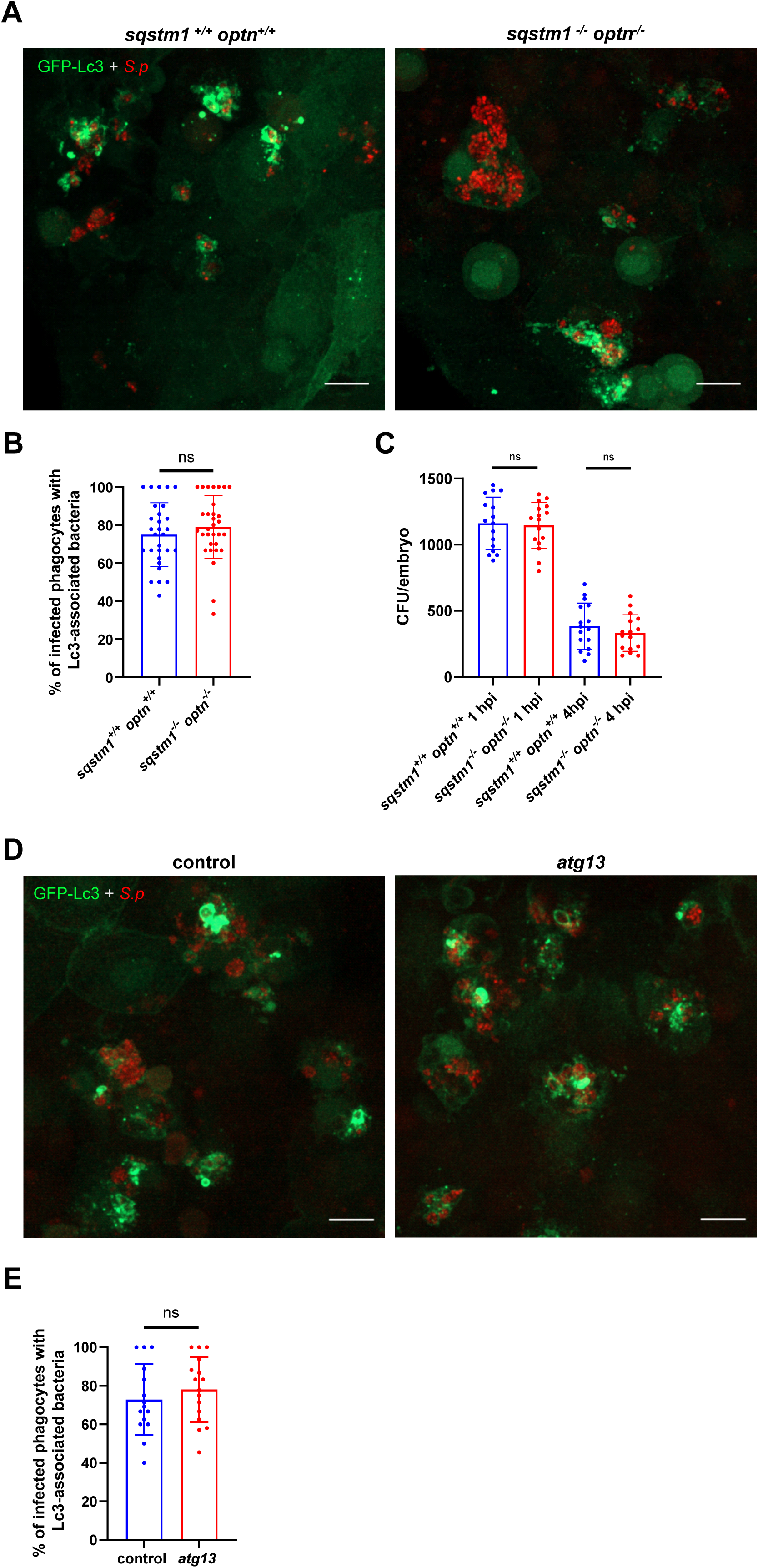
Sqstm1, optineurin and atg13 are not involved in autophagic response to *S. pneumoniae*. (**A**) representative confocal images of *sqstm1*^+/+^/*optn*^+/+^ and *sqstm1*^-/-^ /*optn*^-/-^ double mutant *CMV:GFP-Lc3* larvae infected systemically with approximately 1600 CFU of mKate2-labeled D39 Δ*cps S. pneumoniae* and fixed at 2 hpi Scale bars: 10 µm. (**B**) Quantification of Lc3 associations with intracellular *S. pneumoniae* within infected phagocytes of fixed *CMV:GFP-Lc3 sqstm1*^+/+^/*optn*^+/+^ and *sqstm1*^-/-^/*optn*^-/-^ double mutant larvae. Data are shown as mean ± standard deviation (SD). Data obtained from two independent experiments. (**C**) Numbers of viable colony forming units (CFUs) over time of *sqstm1*^+/+^/*optn*^+/+^ and *sqstm1*^-/-^/*optn*^-/-^ double mutant larvae. Data obtained from two independent experiments. (**D**) representative confocal images of control and *atg13* knockdown (right panel) *CMV:GFP-Lc3* larvae infected systemically with approximately 1600 CFU of mKate2-labeled D39 Δ*cps S. pneumoniae* and fixed at 2 hpi Scale bars: 10 µm. (**E**) Quantification of Lc3 associations with intracellular *S. pneumoniae* within infected phagocytes of fixed *CMV:GFP-Lc3* control and *atg13* knockdown larvae. Data are shown as mean ± standard deviation (SD). n≥15 zebrafish larvae were analyzed.

To exclude potential involvement of the other SLRs such as CALCOCO2 or TAXBP1, we confirmed the lack of xenophagy in pneumococci-infected macrophages by targeting *atg13*, a component of the ULK1 complex that is required for canonical autophagy pathways including xenophagy, but dispensable for LAP [43]. Under conditions of *atg13* CRISPR-mediated knockdown [28], we saw no reduction in Lc3 association with internalized pneumococci in *atg13* knockdown larvae (Fig 5D-E), confirming that the Lc3 response to pneumococci is largely independent of xenophagy.

Overall, these results indicate no apparent role of xenophagy in the autophagic response to pneumococci in zebrafish macrophages suggesting the existence of another non-canonical autophagic mechanism evoked during the *S. pneumoniae* infection that functions alongside LAP and is triggered by pneumolysin.

### A *tecpr1a*-mediated pneumolysin-dependent autophagic pathway occurs in pneumococcus-infected macrophages

In pursuit of defining the hypothesized alternative pneumolysin-dependent non-canonical autophagic route (Fig. 4), we speculated that a recently described CASM pathway called sphingomyelin-TECPR1-induced LC3 lipidation (STIL), could be involved [14]. This response is triggered upon sphingomyelin translocation from the luminal into the cytosolic leaflet of the phagosomal membrane induced by bacterial toxins such as listeriolysin O, from *Listeria monocytogenes*. It has been postulated that cytosolic sphingomyelin exposure is an early indicator of vesicle damage [44]. Importantly, *S. pneumoniae* has been recently shown to induce sphingomyelin exposure to the cytoplasm in phagosomal membranes of infected BMDMs (Shizukuishi et al., 2024). In STIL, the exposed sphingomyelin is recognized by a protein called TECPR1 (tectonin beta-propeller repeat containing 1) to induce single membrane LC3 lipidation [46]. Interestingly, TECPR1 provides E3-ligase-like activity to the ATG12–ATG5 complex, which is essential for the conjugation of LC3 to damaged phagosomes in the absence of ATG16L1 [47]. The zebrafish genome contains two paralogs of *TECPR1* called *tecpr1a* and *tecpr1b* (Fig. 6). We performed an alignment of both zebrafish proteins to the human ortholog and we found profoundly higher amino acid similarity between TECPR1 and Tecpr1a in comparison to Tecpr1b, especially within the ATG5-interacting region (AIR) which appears to be incomplete in Tecpr1b (Fig. 6A-B). In addition, using the Zebrafish Blood Atlas web tool with single cell RNAseq gene expression profile of adult zebrafish hematopoietic cells [48], we found that *tecpr1a*, but not *tecpr1b*, is highly expressed in zebrafish leukocytes (Fig. 6C). Based on this analysis, we hypothesized the Tecpr1a variant to be functional in zebrafish, but nevertheless, we knocked down both zebrafish paralogs, *tecpr1a* and *tecpr1b*, using the CRISPR-Cas9 approach and positively verified their knockdown efficiency (Fig. S2D-E). Interestingly, when challenged by pneumococci, only *tecpr1a* and the *tecpr1a tecpr1b* double knockdown, but not *tecpr1b* knockdown larvae showed partial reduction of Lc3 decoration within infected macrophages (Fig. 6D-E). This result suggests that *tecpr1a* is indeed the functional ortholog of human TECPR1, confirming our bioinformatic analysis (Fig. 6A-C). In order to verify the pneumolysin-dependency of the observed *tecpr1a*-mediated Lc3 decoration, we infected the *tecpr1a* knockdown larvae with the pneumolysin-deficient Δ*ply* strain. We found no additive effect in the reduction of Lc3 associations (Fig. 6F-G), indicating both Tecpr1a and Ply act on the same pathway. To provide further evidence for the involvement of the STIL pathway, we checked if the LC3 response to pneumococci would be independent of ATG16l1, which is not required for STIL [14,47]. Knockdown of *atg16l1* resulted in a significant reduction of Lc3 associations, which was more pronounced than the decrease observed following *cyba* knockdown (Fig. S7D–E), and impaired bacteria clearance (Fig. S7A-C). Although some Lc3 signal remained, these results demonstrate the presence of a largely atg16l1-independent autophagic response to pneumococci in zebrafish macrophages, consistent with the requirements of the STIL pathway.

**Figure 6.**
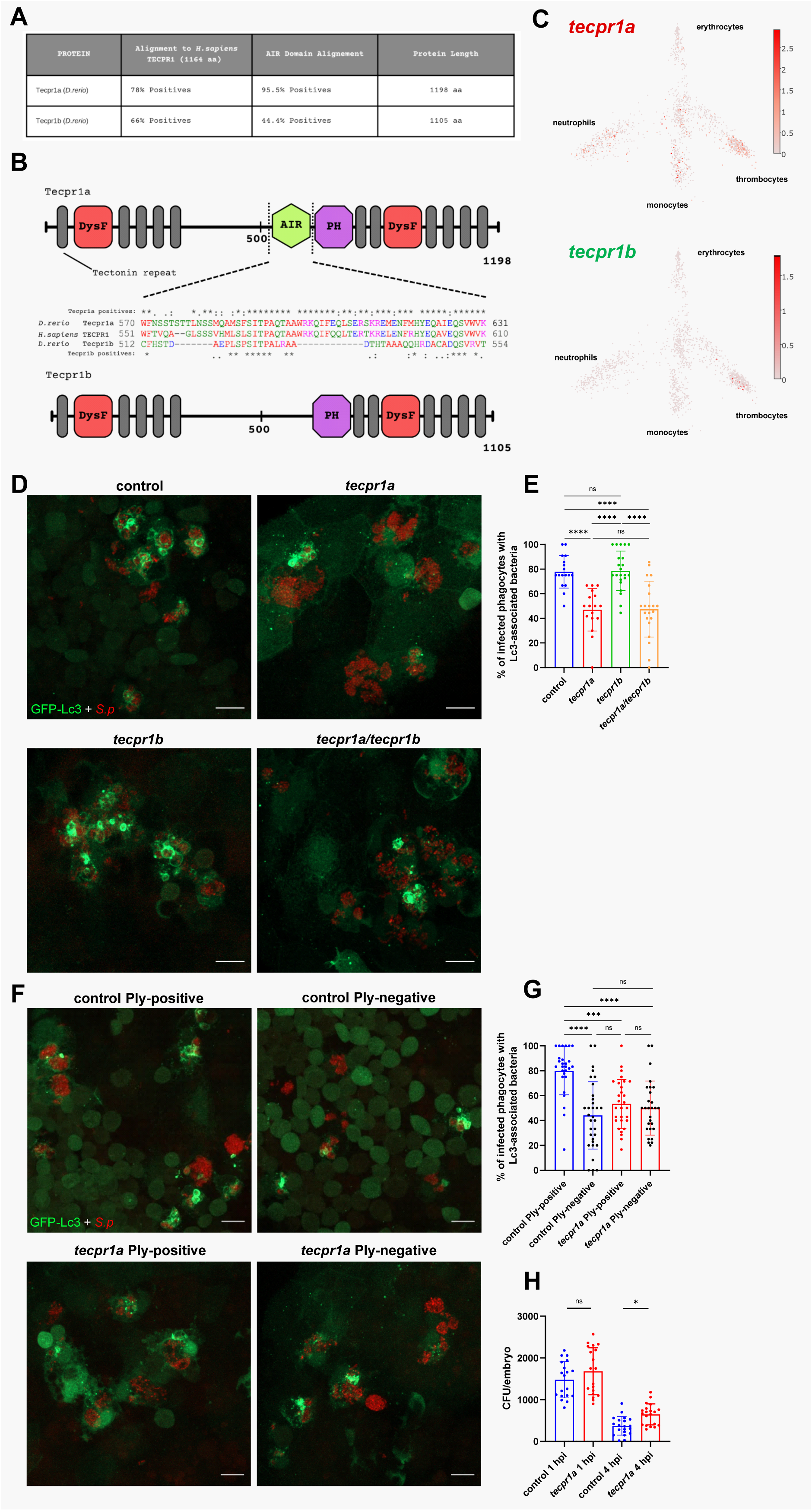
Tecpr1a, a likely ortholog of human TECPR1, controls Ply-mediated autophagic response to pneumococci. (**A**) Table summarizing the alignment of zebrafish Tecpr1a and Tecpr1b proteins to the human TECPR1 protein. Columns display percentage positives (identical amino acids, as well as conserved or semiconserved substitutions) and length of the proteins. Alignments sequence similarities are shown for the full-length protein and the AIR domain, which exhibits the greatest divergence between the zebrafish paralogues (Tecpr1a and Tecpr1b). (**B**) Schematic representation of zebrafish Tecpr1a and Tecpr1b proteins, showing their domains: DysF - Dysferlin, AIR - ATG5-interacting region, and PH - Pleckstrin homology domain. Alignment of the AIR domain amino acid sequence of Human TECPR1 and the zebfafish orthologs, Tecpr1a and Tecpr1b, is shown between the schematics. Symbols denote: (*) identical residues, (:) conserved substitutions, and (.) semiconserved substitutions. (**C**) Gene expression of adult zebrafish leukocytes determined using the Zebrafish Blood Atlas [48]. Each dot represents a separate scRNAseq sample (cell); replicates were performed across multiple zebrafish wild-type and transgenic strains. Each arm of the schematic represents a separate blood cell population (labeled). Deeper color indicates higher expression (log_10_ scale bars described for each gene). (**D**) representative confocal images of control, *tecpr1a*, *tecpr1b* and *tecpr1a*/*b* double knockdown *CMV:GFP-Lc3* larvae infected systemically with approximately 1600 CFU of mKate2-labeled D39 Δ*cps S. pneumoniae* fixed at 2 hpi. (**E**) Quantification of Lc3 associations with intracellular *S. pneumoniae* within infected phagocytes of fixed *tecpr1a, tecpr1b* and *tecpr1a/b* double knockdown *CMV:GFP-Lc3* larvae. Data are shown as mean ± standard deviation (SD). n≥17 zebrafish larvae were analyzed. (**F**) representative confocal images of control or *tecpr1a* knockdown *CMV:GFP-Lc3* larvae infected systemically with approximately 1600 CFU of mKate2-labeled Ply-positive (D39 Δ*cps*) or Ply-negative (D39 Δ*cps* Δ*ply*) *S. pneumoniae* fixed at 2 hpi. (**G**) Quantification of Lc3 associations with intracellular Ply-positive or Ply-negative *S. pneumoniae* within infected phagocytes of control or *tecpr1a* knockdown *CMV:GFP-Lc3* larvae fixed at 2 hpi. Data are shown as mean ± standard deviation (SD). Data obtained from two independent experiments. (**H**) Numbers of viable colony forming units (CFUs) over time of control and *tecpr1a* knockdown larvae. Data obtained from two independent experiments.

Upon discovery of this pathway in pneumococcus-infected macrophages, we set out to determine the role of STIL in pneumococcal killing. The results showed that Tecpr1a-depteted larvae clear the intravenously-administered pneumococci slower than their wild-type siblings. (Fig 6H). Together, these results demonstrate a host-protective role of a non-canonical pathway called STIL that has not been previously been implicated in pneumococcal infection.

### LAP and STIL independently initiate autophagic pathways in response to pneumococci

Our data suggests the occurrence of two simultaneous non-canonical autophagic pathways of different nature in response to pneumococci: LAP and STIL, which are both part of the recently recognized and rapidly expanding family of CASM pathways, where LC3/Atg8 is conjugated to single membranes of phagosomes and other organelles [13,14]. In order to confirm the coexistence of these phenomena we knocked down the two specific factors responsible for each of them: *cyba* for LAP and *tecpr1a* for STIL. When both genes were knocked down we observed a near complete loss of Lc3 associations (Fig 7A-B). Notably, this effect of double *cyba* and *tecpr1a* knockdown was substantially pronounced than the partial reductions of Lc3 decorations that we previously observed in separate knockdowns of *cyba* (Fig 3A-B) or *tecpr1a* (Fig. 6 D-E). This additive effect of *cyba* and *tecpr1a* gene silencing suggests that indeed we observe two separate autophagic pathways evoked in response to internalized pneumococci. Finally, to compare the contributions of LAP and STIL to pneumococcal killing, we determined the level of bacteria in both single and double *cyba tecpr1a* knockdown. This analysis showed that bacteria were killed at a lower pace in larvae where both LAP and STIL were inhibited compared with either of the single knockdown counterparts (Fig. 7C). Collectively, these results confirm the coexistence of LAP and STIL pathways, which both serve host-protective roles and act independently on intracellular pneumococci.

**Figure 7.**
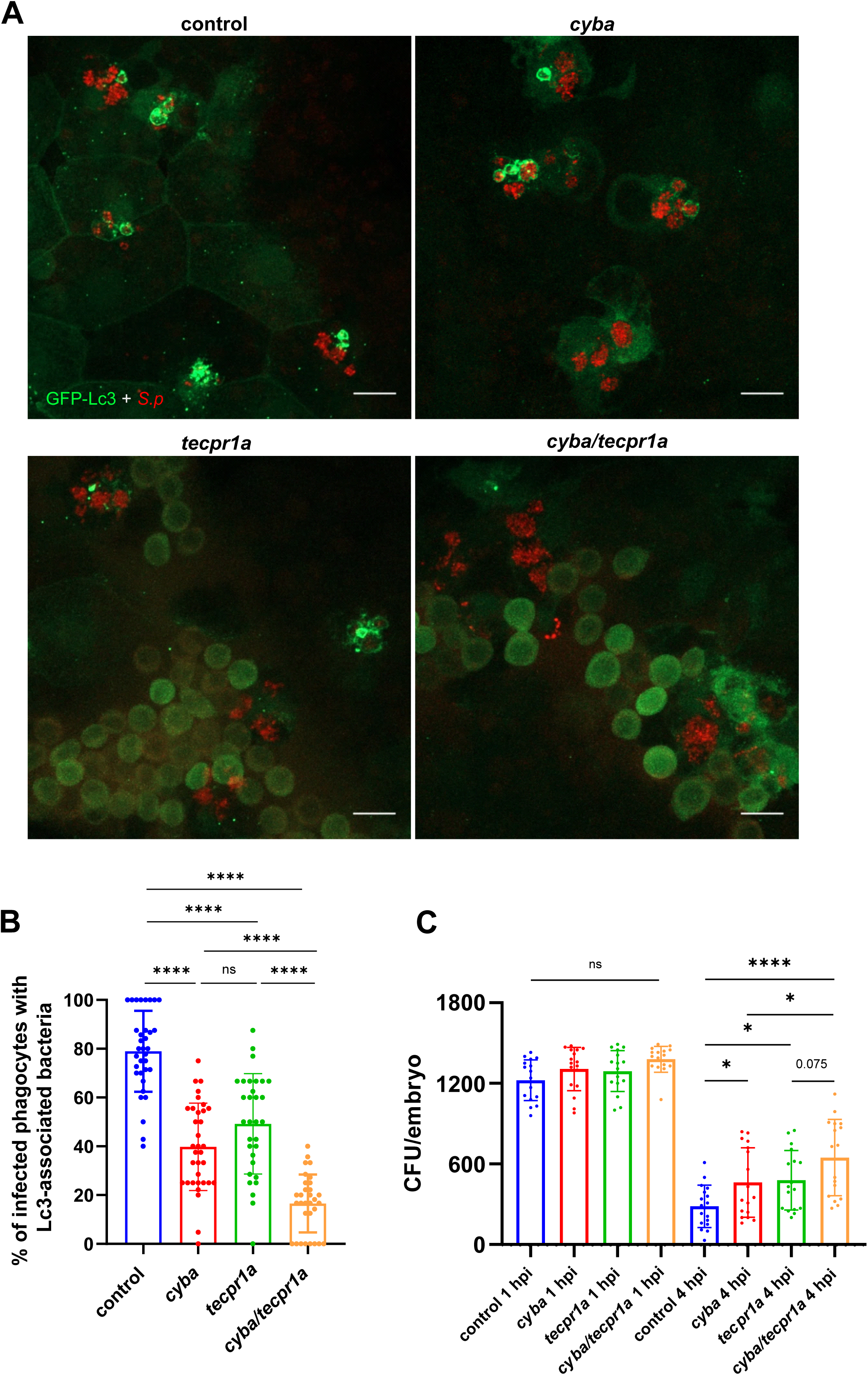
Both LAP and STIL are evoked in response to *S. pneumoniae*. (**A**) representative confocal images of control, *cyba, tecpr1a* and *cyba*/*tecpr1a* double knockdown *CMV:GFP-Lc3* larvae infected systemically with approximately 1600 CFU of mKate2-labeled D39 Δ*cps S. pneumoniae* fixed at 2 hpi. (**B**) Quantification of Lc3 associations with intracellular *S. pneumoniae* within infected phagocytes of fixed *cyba, tecpr1a* and *cyba*/*tecpr1a* double knockdown *CMV:GFP-Lc3* larvae. Data are shown as mean ± standard deviation (SD). Data obtained from two independent experiments. (**C**) Numbers of viable colony forming units (CFUs) over time of control, *cyba*, *tecpr1a and cyba*/*tecpr1a* double knockdown larvae. Data obtained from two independent experiments.

### Another Ply- and Atg16l1-dependent, ROS- and Tecpr1a-independent pathway occurs in response to pneumococcal infection

Although simultaneous knockdown of *cyba* and *tecpr1a* resulted in a strong reduction of Lc3 lipidation, nearly 20% of infected macrophages remained to exhibit Lc3-positive bacteria-containing vesicles. This suggested the existence of a third, distinct autophagic pathway which could be ROS-independent [21]. To investigate this further, a triple knockdown targeting *cyba*, *tecpr1a*, and *atg16l1* was performed. This resulted in a near-complete loss of Lc3 recruitment, reaching level comparable that observed in *atg5*-deficient larvae, were Lc3 lipidation was fully abolished (Fig. 8). Similar to *atg5* CRISPants, the triple knockdown did not affect the survival of larvae upon infection (Fig. S9B). Additionally, to validate the role of pneumolysin in this context, the *cyba tecpr1a* double knockdown larvae were infected with Ply-deficient pneumococci. In this situation, the Lc3 lipidation was also reduced to nearly undetectable levels, indicating that the observed remaining Lc3 signal in the double knockdown background was dependent on this pneumococcal toxin (Fig. 8).

**Figure 8.**
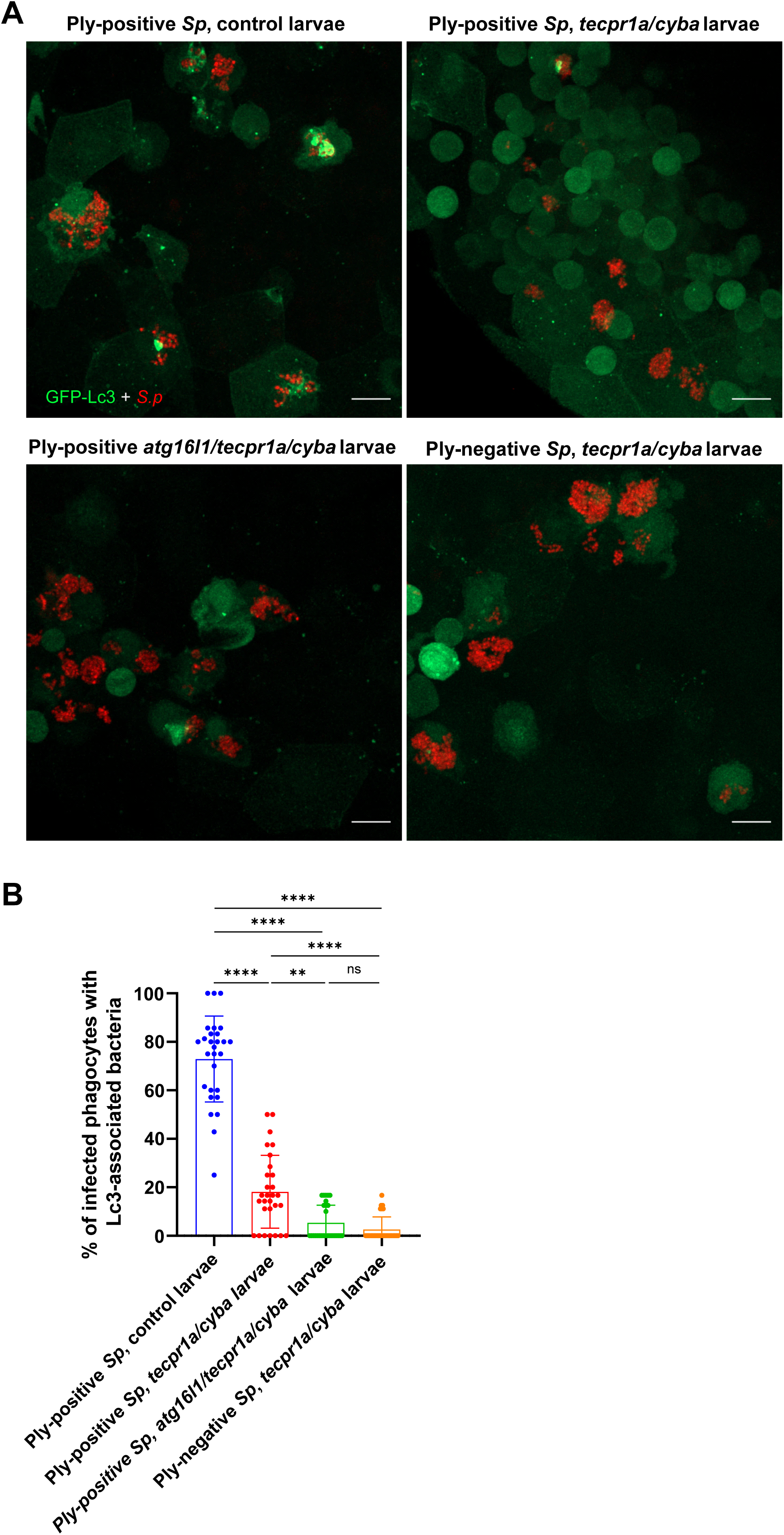
Involvement of ROS-independent LAP-like CASM of *S. pneumoniae*-containing vesicles. (**A**) representative confocal images of control, *tecpr1a/cyba* double knockdown or *atg16l1/tecpr1a/cyba* triple knockdown *CMV:GFP-Lc3* larvae infected systemically with approximately 1600 CFU of Ply-positive (D39 Δ*cps*) and Ply-negative (D39 Δ*cps* Δ*ply*) mKate2-labeled *S. pneumoniae* at 2 hpi. (**B**) Quantification of Lc3 associations with intracellular PLY-positive or PLY-negative *S. pneumoniae* within infected phagocytes of control, *tecpr1a/cyba* double knockdown or *atg16l1/tecpr1a/cyba* triple knockdown *CMV:GFP-Lc3* larvae. Data are shown as mean ± standard deviation (SD). Data obtained from two independent experiments.

All these findings confirmed that the Lc3-positive vesicles observed in the double knockdown derived from a separate, non-canonical autophagy mechanism dependent on both pneumolysin and Atg16l1, but independent of ROS and act in parallel with LAP and STIL, contributing to the Lc3 response in infected macrophages.

## Discussion

Autophagy-related mechanisms, both canonical and non-canonical, are considered to be an important component of host defense against various infections, acting primarily as a protective mechanism, and therefore could be exploited in designing novel therapies [49]. In this study, we aimed to investigate the role of autophagy in the innate immune response to *Streptococcus pneumoniae* systemic infection using a zebrafish larval model. We found that the great majority of infected macrophages contained bacterial clusters surrounded by Lc3 at the early stage of infection (2 hpi) confirming involvement of autophagy in the host response to pneumococci. Importantly, both capsulated and unencapsulated strains, once internalized, were found to effectively activate Lc3 recruitment to pneumococcus-containing vesicles. The LC3 conjugation requires ATG5 which is involved in the covalent attachment of LC3 to the phospholipids of the membrane which is an essential step in the autophagy process [19,31]. By diminishing the autophagic response through knockdown of *atg5* we observed impaired bacterial clearance in our model. This emphasized the significance of autophagy in facilitating the clearance of intracellular pneumococci by zebrafish macrophages and prompted us to investigate which autophagy-related pathway is responsible for this response. We took a combinatory approach by targeting pathogen and host components, both separately and in combination to determine the autophagic pathways present in *S. pneumoniae*-infected macrophages.

The antimicrobial, host-protective function of LC3-associated phagocytosis (LAP) has been reported in several studies, demonstrating its effective role in bacterial degradation [11,50,51]. The significance of this pathway had been recognized well before its inclusion in the recently defined CASM family and it is supported by studies analyzing microbial survival mechanisms evolved to evade or escape this process to prolong their survival [52–54]. Our study demonstrates that LAP occurs in zebrafish macrophages upon internalizing pneumococci. A significant but partial reduction in Lc3 recruitment and slower bacterial clearance induced by knocking down *cyba* or *rubcn*, genes required for LAP, underscores the importance of this pathway in the host defense against pneumococci. Importantly, our data confirms *in vitro*-derived results where LAP also plays a protective role in pneumococci-infected BMDMs [26]. Interestingly, Inomata and colleagues have also reported incomplete inhibition of LC3 recruitment in *Rubcn* or *Cybb/Nox2* knockout macrophages, which additionally prompted us to search for another autophagic pathway present in response to pneumococci. Moreover, in accordance with Inomata and others, we observed significant decrease of Lc3 associations to pneumococci in the absence of pneumolysin and we did not find any evidence of selective autophagy involvement in *S. pneumoniae*-infected macrophages *in vivo*.

Indeed, our results show that Lc3 recruitment triggered by pneumococci upon internalization by macrophages is partially-dependent on bacterial pneumolysin. By targeting components of different autophagy pathways, we were able to match pathogen-derived pneumolysin with particular facets of the autophagic response. Inomata *et al.* posited that pneumolysin induces LAP, however no combinatory approaches with Δ*ply* mutant in LAP-deficient macrophages have been performed [26]. By testing wild-type and Δ*ply* mutant pneumococcal strains against pathway-specific host genes in the zebrafish model, we here show that Ply activates another non-canonical autophagy pathway, which acts independently of ROS and Atg16l1, but is dependent on Tecpr1a.

TECPR1 was initially shown to play a vital role in promoting selective autophagy through interaction with WIPI2 [55]. However, recent studies have revealed an additional function of TECPR1 via its DysF domains [46,56], where this protein recognizes sphingomyelin present on the outer layer of disturbed phagosomal or lysosomal membranes to induce CASM in a process called STIL. Interestingly, this alternative non-canonical pathway challenges the traditional view that ATG16L1 is the sole E3-like enzyme responsible for lipidation of LC3, highlighting the versatility and complexity of autophagic regulation [57].

We have found in zebrafish that a knockdown of *tecpr1a*, a likely ortholog of TECPR1, leads to a partial loss of Lc3 decoration of internalized pneumolysin-positive *S. pneumoniae*. When pneumolysin mutant bacteria were administered, no additional effect of *tecpr1a* depletion was observed suggesting the role of pneumolysin in perturbing the phagosomal membrane and Tecpr1a in phagosomal membrane Lc3 decoration. Importantly, in the *cyba tecpr1a* double knockdown we observed a near-complete loss of Lc3 associations. Moreover, in *cyba* knockdown larvae injected with Δ*ply* pneumococci, no Lc3 decoration was present, indicating the occurrence of two independent, differentially induced non-canonical pathways, namely LAP and STIL. A similar phenomenon was previously observed *in vitro* in BMDMs infected with *L. monocytogenes* evoking two distinct non-canonical autophagy pathways [29]. One of them is LAP and the other ROS-independent, listeriolysin O-mediated, was coined by the authors as pore-forming toxin-induced non-canonical autophagy (PINCA). At the time, the authors did not know the factors responsible for PINCA, therefore they were unable to determine its function in *L. monocytogenes* infection. It is conceivable that PINCA observed by Glushko and colleagues [29] was in fact STIL.

Another similar case of parallel pathways has been observed in MEFs *in vitro* at early stages of infection with pneumolysin-expressing *S. pneumoniae*, where LC3 lipidation occurred independently of RB1CC1/FIP200 (a member of the ULK1 complex required for canonical autophagy) and also independently of ROS leading to LAPosome-like vacuoles formation. However, since the observed process requires ATG16L1 [21], it appears to be another form of CASM, distinct from STIL. Indeed, our data revealed the existence of an equivalent non-canonical autophagy pathway in macrophages, in addition to LAP and STIL. This another CASM depended on both ATG16L1 and pneumolysin, but occurred independently of ROS, matching the mechanism described in non-professional phagocytes [21]. Its identification not only clarifies that Lc3 recruitment was not completely abolished in the absence of LAP and STIL components alone, but also underline the complexity of host defense mechanisms involving non-canonical autophagy.

Interestingly, Ogawa and colleagues have also reported a hierarchical autophagic response in which LAPosomes undergo Sqstm1- and Optn-mediated xenophagy at later stages of infection [21,58]. However, no such phenomenon involving xenophagy factors was observed in macrophages, neither in our study nor in that of others [26]. The various mechanisms evoked by pneumococci infection in different cells are most likely caused by differences between professional and non-professional phagocytes, such as the higher levels of ROS production of macrophages compared to fibroblasts or the degree of phagosomal acidification [26].

The observed differences, particularly the lack of xenophagy activation in macrophages, are most likely caused by differences between professional and non-professional phagocytes, such as the higher levels of ROS production of macrophages compared to fibroblasts or the degree of phagosomal acidification [26].

We propose the following model of the fate of pneumococci within macrophages (Fig. 9). The bacteria are internalized by macrophages and trapped within Lc3-decorated vesicles, which could be formed by three distinct mechanisms depending on the nature of phagosomal membrane perturbation. Florey and others have hypothesized that any process that promotes osmotic imbalances within the endolysosomal system would have the potential to activate LAP-like LC3 lipidation [59]. Therefore, we believe that in our model, the ROS-induced osmotic imbalance in phagosomes containing pneumococci leads to LAP. Alternatively, pneumolysin produced by bacteria trapped in phagosomes inflicts other forms of membrane perturbation, leading to either ROS-independent Lc3 lipidation via unknown mechanism or sphingomyelin presence on the outer side of phagosomal envelope and resulting in STIL. This last scenario is additionally supported by a recent report demonstrating induction of sphingomyelin exposure to the cytoplasm in phagosomal membranes of *S. pneumoniae*-infected BMDMs [60]. Both LAP and STIL enhance pneumococcal killing within zebrafish macrophages as knockdown of pathway-specific components slowed down the inactivation of bacteria *in vivo* but did not completely impair it. This suggests that classical phagosome maturation still occurs in the absence of these pathways and the internalized pneumococci are still trafficked to increasingly acidified vesicles, eventually leading to their degradation.

**Figure 9.**
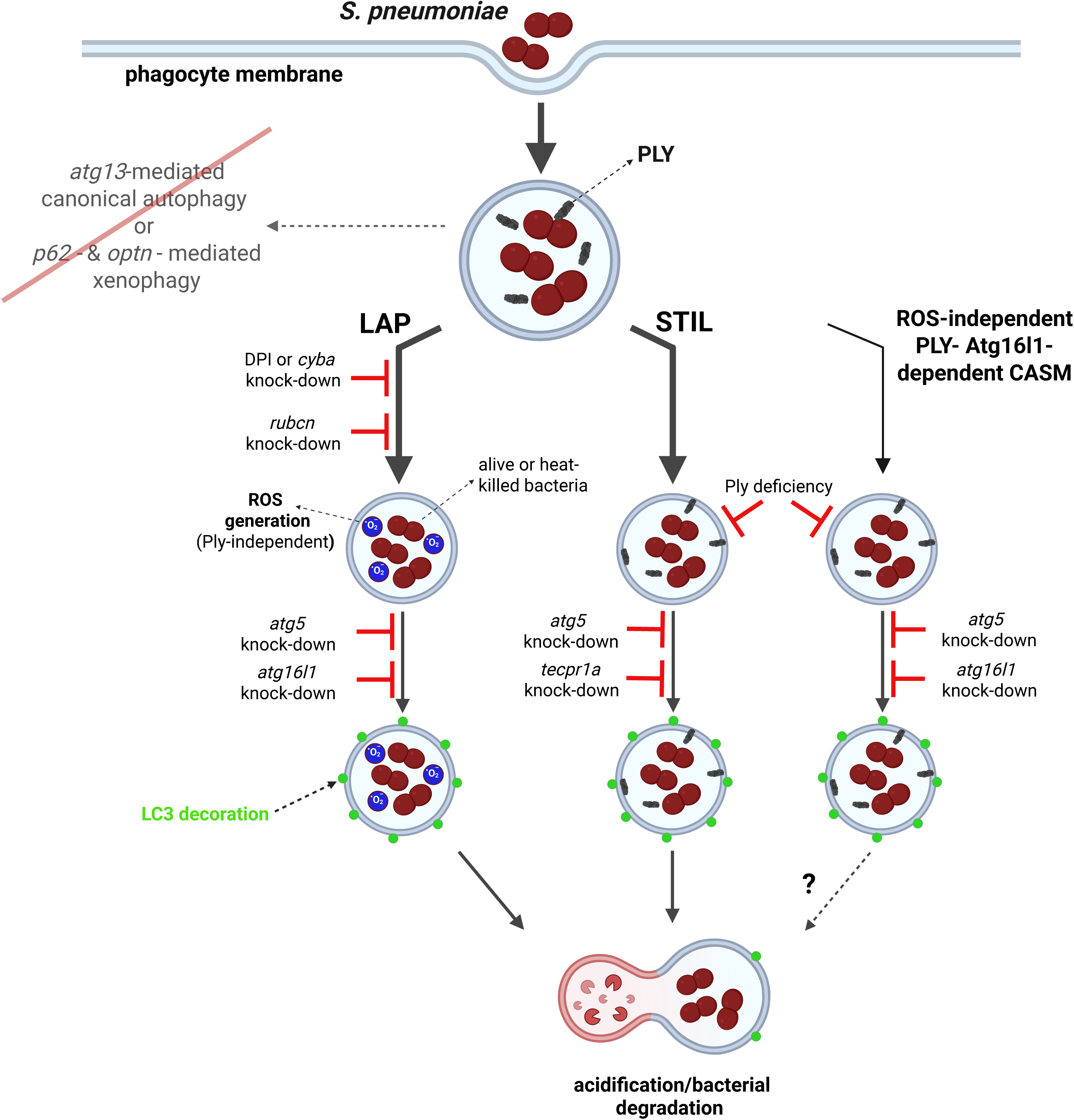
Model of autophagic response to pneumococci within zebrafish macrophages. After internalization by phagocytes, pneumococci are sequestered within Lc3-decorated vacuoles which are formed by three independent mechanisms: ROS-dependent LAP or two ROS-independent Ply-mediated pathways including STIL. LAP occurs as a result of osmotic imbalance within bacteria-containing phagosome caused by ROS production by NADPH oxidase. On the other hand, STIL is induced upon membrane perturbation caused by secretion of bacterial pneumolysin manifested by sphingomyelin presence on the outside leaflet of the phagosome. STIL is dependent on Tecpr1a which recognizes sphingomyelin and mediates Lc3 lipidation. Lastly, the Atg16l1- and Ply-dependent pathway contributes to the smallest portion of Lc3 lipidation. However, the specific mechanisms controlling this process remain to be revealed. Following Lc3-assisted vacuolar sequestration, pneumococci are more efficiently targeted for lysosomal degradation and the Lc3 decoration is removed. The classical xenophagy receptors (sqstm1 and optineurin) as well as the canonical autophagy initiation molecule (atg13) are dispensable in this model. Despite the absence of these xenophagy-mediating factors, the bacteria are efficiently degraded. Figure created with BioRender.

Collectively, the presented results markedly advance our knowledge of the innate immune response to pneumococcal infections. By elucidating the abundant activation of various CASM pathways, and probably negligible role of xenophagy in professional phagocytes our study highlights potential therapeutic targets that could be exploited to enhance host defense mechanisms. Therefore, in addition to LAP, we believe it is important to address other non-canonical autophagy pathways in order to control pneumococcal infection. This may offer novel ways for treatment, especially in cases where traditional antibiotic-based therapies are insufficient due to rising bacterial resistance, opening the door to innovative immunomodulatory approaches that could improve clinical outcomes in pneumococcal infections.

## Materials and Methods

### Ethics statement

All zebrafish experiments were conducted in accordance with the European Community Council Directive 2010/63/EU for the Care and Use of Laboratory Animals of Sept. 22, 2010 (Chapter 1, Article 1 no.3) and National Journal of Law act of Jan. 15, 2015, for Protection of animals used for scientific or educational purposes (Chapter 1, Article 2 no.1). All experiments with zebrafish were performed on embryos/larvae up to 120 h post fertilization and were following ARRIVE guidelines. The Jagiellonian University Zebrafish Core Facility (ZCF) is a licensed breeding and research facility (District Veterinary Inspectorate in Krakow registry; Ministry of Science and Higher Education record no. 022 and 0057).

### Zebrafish husbandry

Adult zebrafish were maintained by Zebrafish Core Facility stac at the Jagiellonian University in accordance with the standard protocol and international guidelines specified by the EU Animal Protective Directive 2010/63/EU. Fish were kept in a continuous recirculating closed system aquarium with a light/dark cycle of 14/10 h at 28°C. Larvae were incubated in standard E3 medium (5 mM NaCl, 0.17 mM KCl, 0.33 mM CaCl_2_, 0.33 mM MgSO_4_) at 28.5°C according to standard protocols [61]. All zebrafish lines used in this study are listed in Table S1.

### Bacterial cultures and infection experiments

All *Streptococcus pneumoniae* strains used in this study were derived from serotype 2 D39 strain [62] and are listed in Table S1. Pneumococci were grown on prepared Tryptic Soy Agar 5% sheep blood plates (TSAII 5% SB; Becton Dickinson [BD], B21261X) at 37°C in an atmosphere containing 5% CO_2_. Liquid cultures were maintained in Todd-Hewitt Broth (BD, 249240) supplemented with 0.5% yeast extract (Gibco, 212750) at 37°C in an atmosphere containing 5% CO_2_. Bacteria were harvested at mid-log phase (OD_600_ between 0.2 - 0.3) and subsequently centrifuged (5000 g, 10 min) and immediately resuspended in phosphate-bucered saline (Bioshop, PBS404.100) to obtain the desired concentration of approximately 1600 CFU/nl. Such prepared bacteria were microinjected intravenously into 36 hpf zebrafish larvae as described previously [63]. Briefly, larvae were anaesthetized with 0.02% buffered tricaine (Sigma-Aldrich, A5040) and then transferred onto 3% w:v methylcellulose (Sigma-Aldrich, M7027) and injected individually with 1 nl of bacteria inoculum using microcapillary pipettes.

### Staining of bacteria with pHrodo-Red and Alexa Fluor 647 S-ester dyes

The pHrodo Red (ThermoFisher, P36600) and Alexa Fluor 647 (ThermoFisher, A37573) succinimidyl-ester dyes were dissolved in DMSO (Sigma-Aldrich, D8418) to the final concentrations of 2.5 mM and 8 mM, respectively. In the indicated experiments, *S. pneumoniae* strains were stained as previously described [25].

### Generation of ***Δ****ply* mutant in S. pneumoniae

The *S. pneumoniae* Δ*ply* deletion strain was generated using linear PCR fragments. Briefly, two ∼1-kb flanking regions of each gene were amplified and an antibiotic resistance marker placed between them using isothermal assembly. Assembled PCR products were transformed directly into *S. pneumoniae*. In all cases, deletion primers were given the typical name: “gene-designation”_5FLANK_F/R for 5′ regions and “gene-designation”_3FLANK_F/R for 3′ regions, antibiotic markers were amplified from Δ*bgaA* strains [64] using the AntibioticMarker_F/R primers. Successful gene deletion strains were confirmed using diagnostic PCR. All bacterial strains and oligonucleotides used in this study are listed in Table S2.

### Morpholino- and CRISPR/Cas9-Mediated Knockdown

The *cyba* morpholino oligonucleotides [32] (Gene Tools, 5’-ATCATAGCATGTAAGGATACATCCC-3’) were dissolved in MilliQ water to obtain the required concentrations of 1 mM. The 1 nl volume of morpholino was injected into the yolk of 1–2 cell stage zebrafish embryos using a microinjector. Standard control morpholino (Gene Tools, 5’-CCTCTTACCTCAGTTACAATTTATA-3’) was used as a negative control. For CRISPR-Cas9-mediated *atg16l1, rubcn*, *tecpr1a* and *tecpr1b* knockdown, the online web tool CHOPCHOP was used to design a gene-specific guide CRISPR of the following sequences listed in Table S3. The synthetic guide RNA consisting of gene-specific CRISPR RNA (crRNA, Merck, VC40002) and transactivating RNAs (tracrRNA, Merck, TRACRRNA05N) in combination with Cas9 nuclease protein (NEB, M0386M) was used for gene editing. TracrRNA and crRNA were resuspended to a concentration of 50 μM in nuclease-free water. The gRNA-Cas9 complexes were assembled before injections using a 1:1:1 ratio of crRNA:tracrRNA:Cas9 nuclease with the final concentrations of 16.6 μM, 16.6 μM, and 6.6 μM, respectively. As controls, only Cas9 + tracrRNA were injected without crRNA. Knockdown verification was performed by the PCR use of primers with the sequences presented in the Table S2. For *atg5* knockdown, a previously published single guide RNA (sgRNA, Merck, VC40003) was used. Briefly, 1 μl 25 μM sgRNA was gently mixed with 1 μl 25 μM Cas9 and 3 μl H2O with the final concentration of 5 μM and 2 nl was injected. The mutagenesis efficacy for *atg5* was determined by restriction fragment length polymorphism analysis using the MslI enzyme as previously described [65].

### DPI treatment

At 1 h before infection, larvae were bath-treated with 100 µM DPI (Sigma-Aldrich, D2926) in E3 medium. The larvae were then infected and kept in DPI solution until fixation. DMSO at 0.1% was used as vehicle control.

### Immunostaining

Larvae were fixed with 4% paraformaldehyde(PFA, ThermoFisher, AAJ19943K2) supplemented with 0.5% Triton X-100 (Sigma-Aldrich, X-100) in PBS and incubated overnight at 4°C. Subsequently, fixed fish were washed four times for 20 min in PBS with 0.5% Triton X-100 and 1% DMSO (PBS-DTx). Next, the larvae were digested in 10 μg/ml proteinase K (Sigma-Aldrich, 1073930010) for 10 min at 37°C. Then, larvae were quickly washed three timed for 5 minutes and blocked with PBS-DTx containing 1% bovine serum albumin (BSA; Sigma-Aldrich, A9418) for 1h at room temperature and incubated overnight at 4°C in mono-and polyubiquitinated conjugates recombinant mouse monoclonal antibody (Enzo Life Siences, UBCJ2) diluted in the blocking buffer 1:250. Next, larvae were washed in blocking solution four times for 20 minutes and incubated at room temperature for 2 h in blocking solution of secondary goat anti-mouse Alexa Fluor 405 antibody (Invitrogen, A-31553) in 1:300 followed with four times washes in PBS with 0.5% Triton-X100.

### Microscopy Imaging and Analysis

Live anaesthetized or paraformaldehyde-fixed larvae were immersed in 1% (w/v) low-melting-point agarose solution in E3 medium and mounted laterally on a dish with a glass bottom. Images of the common cardinal vein (top of Fig.1) were acquired using a Zeiss LSM900 Airyscan 2 confocal laser scanning microscope with ZEISS ZEN 3.3 software using the Plan-Apochromat 20x/0.8, C-Apochromat 40x/NA 1.2 W or C-Apochromat 63x/1.20 W objectives. For quantification of the autophagic response within infected phagocytes, for each larva, a total number of observable infected phagocytes were manually determined through the z-stacks of images. Among these total observable infected phagocytes number of infected phagocytes with GFP-Lc3 signals were enumerated, and the percentage of Lc3-positive phagocytes over total observable phagocytes was determined for each larva. Maximum projections were used for representative images. No non-linear normalizations were performed. For quantification of acidification, the pHrodo Red fluorescence intensity, ZEISS ZEN 3.3 software was used. Quantification of bacterial internalization was performed as previously described [66].

### Determination of In Vivo Bacterial Counts

At various times postinfection, approximately 8-12 anesthetized zebrafish larvae were individually transferred with 100cμl of E3 medium into 0.5-ml Screw cap micro tubes (Sarstedt, 72.730.100) containing 2-mm Zirconia beads (Biospec, 11079124ZX) and homogenized using a FastPrep-24 5G homogenizer (MP Biomedicals, 6005500). The homogenates were spotted on TSA plates containing 5% defibrinated sheep blood to determine pneumococcal CFU numbers. The limit of detection was 10cCFU per larva.

### Western Blot

For each immunoblot, 10 μl of each whole-cell lysate was run on a 12% SDS PAGE gel (40 mA, ≈55 min) and transferred onto PVDF membranes (80 min, 50 V). Membranes were blocked with 5% skimmed milk in 1x PBS 0.5% Tween 20 (Sigma-Aldrich, 11332465001) for 1 h (gently rocking, room temperature). Membranes were washed three times with 1x PBS, 0.05% Tween 20, and incubated with primary antibody overnight gently rocking (4°C, 3% BSA in 1x PBS, 0.05% Tween 20, 0.02% sodium azide). For detection of Ply, a mouse anti-Ply antibody was used at a final concentration of 1:1000 (mouse mAb to pneumolysin [PLY-4]; Abcam, AB71810). The blocked PVDF membranes were washed three times with 1x PBS, 0.05% Tween 20. The secondary antibody (goat anti-mouse IgG conjugated to horseradish peroxidase [Bio-Rad, 172-1011]) was used at 1:5000 and applied in 1x PBS with 3% BSA for 1 h at room temperature with gentle shaking. Resulting blots were washed four times in 1x PBS, 0.05% Tween 20 and visualized using the Clarity Western ECL Substrate kit (Bio-Rad, 170-5061) according to the manufacturer’s instructions. Blots were imaged using a Syngene G:BOX Chemi XX9 image quantification device combined with the GeneSys software (v1.6.7.0) using the chemiluminescent detection tools.

### Statistics

GraphPad Prism 10 was used for statistical analysis. Quantifications of percent Lc3-positive phagocytes, CFU counts, bacterial uptake, or pHrodo Red and GFP fluorescence intensity were determined for significance with unpaired parametric t-test for 2 groups and with ANOVA for multiple groups, with Šidák correction for multiple comparisons. Survival curves were analyzed with Log rank (Mantel-Cox) test. Statistical significance was assumed at *P*-values below 0.05.

## Acknowledgements

This work was supported by a National Science Centre of Poland under Sonata Bis 9 project (Grant number: 2019/34/E/NZ6/00137) awarded to T.K.P. and a Medical Research Council (MRC) New Investigator Research Grant (Grant number: MR/S009280/1) awarded to A.K.F.

T.K.P. was funded by Polish National Agency for Academic Exchange under Polish Returns 2019 project (Grant number: PPN/PPO/2019/1/00029/U/0001) and T.C.S. was funded by MRC DiMeN DTP studentship.

The establishment of a Zebrafish Core Facility at the Department of Evolutionary Immunology has been supported by a grant from the Priority Research Area BioS under the Strategic Program Excellence Initiative at Jagiellonian University.

## ABBREVIATIONS

ATG: autophagy related
BMDM: bone marrow-derived macrophage
CASM: conjugation of ATG8 to single membranes
CFU: colony-forming units
Cyba: cytochrome b-245, alpha polypeptide
DPI: diphenyleneiodonium
GFP: green fluorescent protein
hpf: hours post-fertilization
hpi: hours post-infection
LAP: LC3-associated phagocytosis
Map1lc3/Lc3: microtubule-associated protein 1 light chain 3
MEF: mouse embryonic fibroblast
NADPH: nicotinamide adenine dinucleotide phosphate
Optn: optineurin
Ply: pneumolysin
ROS: reactive oxygen species
SLR: sequestosome-like receptors
Sqstm1: sequestosome 1
STIL: sphingomyelin-TECPR1-induced LC3 lipidation
Tecpr1: tectonin beta-propeller repeat containing 1

## References

[1] Weiser JN, Ferreira DM, Paton JC. Streptococcus pneumoniae: Transmission, colonization and invasion. Nat Rev Microbiol 2018;16:355–67. 10.1038/s41579-018-0001-8.

[2] Ikuta KS, Swetschinski LR, Aguilar GR, et al. Global mortality associated with 33 bacterial pathogens in 2019: a systematic analysis for the Global Burden of Disease Study 2019. The Lancet 2022;400:2221–48. 10.1016/S0140-6736(22)02185-7.

[3] van der Poll T, Opal SM. Pathogenesis, treatment, and prevention of pneumococcal pneumonia. The Lancet 2009;374:1543–56. 10.1016/S0140-6736(09)61114-4.

[4] Croucher NJ, Harris SR, Fraser C, et al. Rapid pneumococcal evolution in response to clinical interventions. Science (1979) 2011;331:430–4. 10.1126/science.1198545.

[5] Dockrell DH, Marriott HM, Prince LR, et al. Alveolar Macrophage Apoptosis Contributes to Pneumococcal Clearance in a Resolving Model of Pulmonary Infection. The Journal of Immunology 2003;171:5380–8. 10.4049/jimmunol.171.10.5380.

[6] Flannagan RS, Jaumouillé V, Grinstein S. The cell biology of phagocytosis. Annual Review of Pathology: Mechanisms of Disease 2012;7:61–98. 10.1146/annurev-pathol-011811-132445.

[7] Maxson ME, Grinstein S. The vacuolar-type H+-ATPase at a glance - more than a proton pump. J Cell Sci 2014;127:4987–93. 10.1242/jcs.158550.

[8] Westman J, Grinstein S. Determinants of Phagosomal pH During Host-Pathogen Interactions. Front Cell Dev Biol 2021;8. 10.3389/fcell.2020.624958.

[9] Prajsnar TK, Michno BJ, Pooranachandran N, et al. Phagosomal Acidification Is Required to Kill Streptococcus pneumoniae in a Zebrafish Model. Cell Microbiol 2022;2022. 10.1155/2022/9429516.

[10] Kaufmann SHE, Dorhoi A. Molecular Determinants in Phagocyte-Bacteria Interactions. Immunity 2016;44:476–91. 10.1016/j.immuni.2016.02.014.

[11] Yuan J, Zhang Q, Chen S, et al. LC3-Associated Phagocytosis in Bacterial Infection. Pathogens 2022;11. 10.3390/pathogens11080863.

[12] Sharma V, Verma S, Seranova E, et al. Selective autophagy and xenophagy in infection and disease. Front Cell Dev Biol 2018;6. 10.3389/fcell.2018.00147.

[13] Durgan J, Florey O. Many roads lead to CASM: Diverse stimuli of noncanonical autophagy share a unifying molecular mechanism. Sci Adv 2022;8. 10.1126/sciadv.abo1274.

[14] Figueras-Novoa C, Timimi L, Marcassa E, et al. Conjugation of ATG8s to single membranes at a glance. J Cell Sci 2024;137. 10.1242/jcs.261031.

[15] Shaid S, Brandts CH, Serve H, et al. Ubiquitination and selective autophagy. Cell Death Differ 2013;20:21–30. 10.1038/cdd.2012.72.

[16] Thurston TLM, Wandel MP, Von Muhlinen N, et al. Galectin 8 targets damaged vesicles for autophagy to defend cells against bacterial invasion. Nature 2012;482:414–8. 10.1038/nature10744.

[17] Galluzzi L, Baehrecke EH, Ballabio A, et al. Molecular definitions of autophagy and related processes. EMBO J 2017;36:1811–36. 10.15252/embj.201796697.

[18] Herhaus L, Dikic I. Regulation of Salmonella-host cell interactions via the ubiquitin system. International Journal of Medical Microbiology 2018;308:176–84. 10.1016/j.ijmm.2017.11.003.

[19] Melia TJ, Lystad AH, Simonsen A. Autophagosome biogenesis: From membrane growth to closure. Journal of Cell Biology 2020;219. 10.1083/JCB.202002085.

[20] Ogawa M, Matsuda R, Takada N, et al. Molecular mechanisms of Streptococcus pneumoniae-targeted autophagy via pneumolysin, Golgi-resident Rab41, and Nedd4-1-mediated K63-linked ubiquitination. Cell Microbiol 2018;20. 10.1111/cmi.12846.

[21] Ogawa M, Takada N, Shizukuishi S, et al. Streptococcus pneumoniae triggers hierarchical autophagy through reprogramming of LAPosome-like vesicles via NDP52-delocalization. Commun Biol 2020;3. 10.1038/s42003-020-0753-3.

[22] Yang CS, Lee JS, Rodgers M, et al. Autophagy protein rubicon mediates phagocytic NADPH oxidase activation in response to microbial infection or TLR stimulation. Cell Host Microbe 2012;11:264–76. 10.1016/j.chom.2012.01.018.

[23] Hooper KM, Jacquin E, Li T, et al. V-ATPase is a universal regulator of LC3-associated phagocytosis and non-canonical autophagy. Journal of Cell Biology 2022;221. 10.1083/jcb.202105112.

[24] Masud S, Prajsnar TK, Torraca V, et al. Macrophages target Salmonella by Lc3-associated phagocytosis in a systemic infection model. Autophagy 2019;15:796–812. 10.1080/15548627.2019.1569297.

[25] Prajsnar TK, Serba JJ, Dekker BM, et al. The autophagic response to Staphylococcus aureus provides an intracellular niche in neutrophils. Autophagy 2021;17:888–902. 10.1080/15548627.2020.1739443.

[26] Inomata M, Xu S, Chandra P, et al. Macrophage LC3-associated phagocytosis is an immune defense against Streptococcus pneumoniae that diminishes with host aging. Proc Natl Acad Sci U S A 2020;117. 10.1073/PNAS.2015368117.

[27] He C, Bartholomew CR, Zhou W, et al. Assaying autophagic activity in transgenic GFP-Lc3 and GFP-Gabarap zebrafish embryos. Autophagy 2009;5:520–6. 10.4161/auto.5.4.7768.

[28] Forn-Cuní G, Welvaarts L, Stel FM, et al. Stimulating the autophagic-lysosomal axis enhances host defense against fungal infection in a zebrafish model of invasive Aspergillosis. Autophagy 2023;19:324–37. 10.1080/15548627.2022.2090727.

[29] Gluschko A, Farid A, Herb M, et al. Macrophages target Listeria monocytogenes by two discrete non-canonical autophagy pathways. Autophagy 2022;18:1090–107. 10.1080/15548627.2021.1969765.

[30] Bernut A, Herrmann JL, Kissa K, et al. Mycobacterium abscessus cording prevents phagocytosis and promotes abscess formation. Proc Natl Acad Sci U S A 2014;111. 10.1073/pnas.1321390111.

[31] Dikic I, Elazar Z. Mechanism and medical implications of mammalian autophagy. Nat Rev Mol Cell Biol 2018;19:349–64. 10.1038/s41580-018-0003-4.

[32] Niethammer P, Grabher C, Look AT, et al. A tissue-scale gradient of hydrogen peroxide mediates rapid wound detection in zebrafish. Nature 2009;459:996–9. 10.1038/nature08119.

[33] Masud S, van der Burg L, Storm L, et al. Rubicon-Dependent Lc3 Recruitment to Salmonella-Containing Phagosomes Is a Host Defense Mechanism Triggered Independently From Major Bacterial Virulence Factors. Front Cell Infect Microbiol 2019;9. 10.3389/fcimb.2019.00279.

[34] Periselneris J, Turner CT, Ercoli G, et al. Pneumolysin suppresses the initial macrophage pro-inflammatory response to Streptococcus pneumoniae. Immunology 2022;167:413–27. 10.1111/imm.13546.

[35] Ogryzko N V., Lewis A, Wilson HL, et al. Hif-1α–Induced Expression of Il-1β Protects against Mycobacterial Infection in Zebrafish. The Journal of Immunology 2019;202:494–502. 10.4049/JIMMUNOL.1801139.

[36] Marjoram L, Alvers A, Deerhake ME, et al. Epigenetic control of intestinal barrier function and inflammation in zebrafish. Proc Natl Acad Sci U S A 2015;112:2770–5. 10.1073/PNAS.1424089112/SUPPL_FILE/PNAS.201424089SI.PDF.

[37] Manzanillo PS, Ayres JS, Watson RO, et al. The ubiquitin ligase parkin mediates resistance to intracellular pathogens. Nature 2013;501:512–6. 10.1038/nature12566.

[38] Polajnar M, Dietz MS, Heilemann M, et al. Expanding the host cell ubiquitylation machinery targeting cytosolic Salmonella . EMBO Rep 2017;18:1572–85. 10.15252/embr.201643851.

[39] Zhang R, Varela M, Vallentgoed W, et al. The selective autophagy receptors Optineurin and p62 are both required for zebrafish host resistance to mycobacterial infection. PLoS Pathog 2019;15. 10.1371/journal.ppat.1007329.

[40] Bell SL, Lopez KL, Cox JS, et al. Galectin-8 senses phagosomal damage and recruits selective autophagy adapter tax1bp1 to control mycobacterium tuberculosis infection in macrophages. MBio 2021;12. 10.1128/mBio.01871-20.

[41] Xie J, Meijer AH. Xenophagy receptors Optn and p62 and autophagy modulator Dram1 independently promote the zebrafish host defense against Mycobacterium marinum. Front Cell Infect Microbiol 2023;13. 10.3389/fcimb.2023.1331818.

[42] Gubas A, Dikic I. A guide to the regulation of selective autophagy receptors. FEBS Journal 2022;289:75–89. 10.1111/febs.15824.

[43] Zachari M, Ganley IG. The mammalian ULK1 complex and autophagy initiation. Essays Biochem 2017;61:585. 10.1042/EBC20170021.

[44] Ellison CJ, Kukulski W, Boyle KB, et al. Transbilayer Movement of Sphingomyelin Precedes Catastrophic Breakage of Enterobacteria-Containing Vacuoles. Current Biology 2020;30:2974–2983.e6. 10.1016/j.cub.2020.05.083.

[45] Shizukuishi S, Ogawa M, Kuroda E, et al. Pneumococcal sialidase promotes bacterial survival by fine-tuning of pneumolysin-mediated membrane disruption. Cell Rep 2024;43:113962. 10.1016/j.celrep.2024.113962.

[46] Boyle KB, Ellison CJ, Elliott PR, et al. TECPR1 conjugates LC3 to damaged endomembranes upon detection of sphingomyelin exposure . EMBO J 2023;42. 10.15252/embj.2022113012.

[47] Wang Y, Jefferson M, Whelband M, et al. TECPR1 provides E3-ligase like activity to the ATG5-ATG12 complex to conjugate LC3/ATG8 to damaged lysosomes. BioRxiv 2023:2023.06.24.546289. 10.1101/2023.06.24.546289.

[48] Athanasiadis EI, Botthof JG, Andres H, et al. Single-cell RNA-sequencing uncovers transcriptional states and fate decisions in haematopoiesis. Nat Commun 2017;8. 10.1038/s41467-017-02305-6.

[49] Kimmey JM, Stallings CL. Bacterial Pathogens versus Autophagy: Implications for Therapeutic Interventions. Trends Mol Med 2016;22:1060–76. 10.1016/j.molmed.2016.10.008.

[50] Herb M, Gluschko A, Schramm M. LC3-associated phagocytosis - The highway to hell for phagocytosed microbes. Semin Cell Dev Biol 2020;101:68–76. 10.1016/j.semcdb.2019.04.016.

[51] Grijmans BJM, van der Kooij SB, Varela M, et al. LAPped in Proof: LC3-Associated Phagocytosis and the Arms Race Against Bacterial Pathogens. Front Cell Infect Microbiol 2022;11. 10.3389/fcimb.2021.809121.

[52] Hubber A, Kubori T, Coban C, et al. Bacterial secretion system skews the fate of Legionella-containing vacuoles towards LC3-associated phagocytosis. Sci Rep 2017;7. 10.1038/srep44795.

[53] Köster S, Upadhyay S, Chandra P, et al. Mycobacterium tuberculosis is protected from NADPH oxidase and LC3-associated phagocytosis by the LCP protein CpsA. Proc Natl Acad Sci U S A 2017;114:E8711–20. 10.1073/pnas.1707792114.

[54] Mitchell G, Cheng MI, Chen C, et al. Listeria monocytogenes triggers noncanonical autophagy upon phagocytosis, but avoids subsequent growth-restricting xenophagy. Proc Natl Acad Sci U S A 2017;115:E210–7. 10.1073/pnas.1716055115.

[55] Ogawa M, Yoshikawa Y, Kobayashi T, et al. A Tecpr1-dependent selective autophagy pathway targets bacterial pathogens. Cell Host Microbe 2011;9:376–89. 10.1016/j.chom.2011.04.010.

[56] Kaur N, de la Ballina LR, Haukaas HS, et al. TECPR1 is activated by damage-induced sphingomyelin exposure to mediate noncanonical autophagy . EMBO J 2023;42. 10.15252/embj.2022113105.

[57] Corkery DP, Castro-Gonzalez S, Knyazeva A, et al. An ATG12-ATG5-TECPR1 E3-like complex regulates unconventional LC3 lipidation at damaged lysosomes. EMBO Rep 2023;24. 10.15252/embr.202356841.

[58] Sakuma C, Shizukuishi S, Ogawa M, et al. Individual Atg8 paralogs and a bacterial metabolite sequentially promote hierarchical CASM-xenophagy induction and transition. Cell Rep 2024;43. 10.1016/j.celrep.2024.114131.

[59] Florey O, Gammoh N, Kim SE, et al. V-ATPase and osmotic imbalances activate endolysosomal LC3 lipidation. Autophagy 2015;11:88–99. 10.4161/15548627.2014.984277.

[60] Shizukuishi S, Ogawa M, Akeda Y. Individual Atg8 paralogs exhibit unique properties in streptococcus pneumoniae-induced hierarchical autophagy. Autophagy 2024. 10.1080/15548627.2024.2375707.

[61] Nusslein-Volhard C, Dahm R. Zebrafish: A Practical Approach. Oxford University Press; Illustrated Edition (November 21, 2002); 2002.

[62] Lanie JA, Ng WL, Kazmierczak KM, et al. Genome sequence of Avery’s virulent serotype 2 strain D39 of Streptococcus pneumoniae and comparison with that of unencapsulated laboratory strain R6. J Bacteriol 2007;189:38–51. 10.1128/JB.01148-06.

[63] Prajsnar TK, Cunliffe VT, Foster SJ, et al. A novel vertebrate model of Staphylococcus aureus infection reveals phagocyte-dependent resistance of zebrafish to non-host specialized pathogens. Cell Microbiol 2008;10:2312–25. 10.1111/j.1462-5822.2008.01213.x.

[64] Fenton AK, Mortaji L El, Lau DTC, et al. CozE is a member of the MreCD complex that directs cell elongation in Streptococcus pneumoniae. Nat Microbiol 2016;2. 10.1038/nmicrobiol.2016.237.

[65] Chen XK, Yi ZN, Lau JJY, et al. Distinct roles of core autophagy-related genes in zebrafish definitive hematopoiesis. Autophagy 2023;20:830–46. 10.1080/15548627.2023.2274251;SUBPAGE:STRING:FULL.

[66] Salamaga B, Prajsnar TK, Jareño-Martinez A, et al. Bacterial size matters: Multiple mechanisms controlling septum cleavage and diplococcus formation are critical for the virulence of the opportunistic pathogen Enterococcus faecalis. PLoS Pathog 2017;13:e1006526. 10.1371/JOURNAL.PPAT.1006526.

