## Supplementary Material for "Pneumolysin-dependent and independent non-canonical autophagy processes mediate host defense against pneumococcal infection"

**Table S1.** Zebrafish lines used in this study.

| **Zebrafish line name** | **Description** | **Reference** |
| --- | --- | --- |
| AB/TL | Wild-type strain resulted from crossing AB and Tupfel long fin strains | European Zebrafish Resource Centre (EZRC) |
| *Tg*(*CMV*:*GFP-map1lc3b*)*^zf155^* | GFP reporter transgenic zebrafish  for Lc3 | [27] |
| *Tg(mpeg1:mCherryF)^ump2^* | Membrane-localized mCherry reporter for zebrafish macrophages | [30] |
| *optn ^ibl51/ibl51^* / GFP-Lc3 | *optn^ibl51^* mutant line carrying a transgenic GFP-Lc3 reporter | [39] |
| *optn*^+/+^/GFP-Lc3 | siblings of *optn^ibl51/ibl51^* /GFP-Lc3 carrying a transgenic GFP-Lc3 reporter | [39] |
| *sqstm1 ^ibl52/ibl52^* /GFP-Lc3 | *sqstm1^ibl52^* mutant line carrying a transgenic GFP-Lc3 reporter | [39] |
| *sqstm1*^+/+^/GFP-Lc3 | siblings of *sqstm1*^i^*^bl52/ibl52^*/GFP-Lc3 carrying a transgenic GFP-Lc3 reporter | [39] |
| *optn ^ibl51/ibl51^/sqstm1 ^ibl52/ibl52^*/GFP-Lc3 | *optn ^ibl51/ibl51^/sqstm1 ^ibl52/ibl52^* double mutant line carrying a transgenic GFP-Lc3 reporter | [41] |
| *optn*^+/+^/*sqstm1*^+/+^/GFP-Lc3 | siblings of *optn ^ibl51/ibl51^/sqstm1 ^ibl52/ibl52^*/GFP-Lc3 carrying a transgenic GFP-Lc3 reporter | [41] |
| *TgBAC(il1b:GFP)^sh445^* | GFP reporter transgenic zebrafish  for *il1b* (interleukin 1, beta) expression | [35] |
| *TgBAC(tnfa:GFP)^pd1028^* | GFP reporter transgenic zebrafish  for *tnfa* (tumor necrosis factor a) expression | [36] |

**Table S2.** Bacterial strains and oligonucleotides for *∆ply generation* used in this study.

| **Bacterial strain name** | **Description** | **Reference** |
| --- | --- | --- |
| D39 *∆cps* | Unencapsulated *S. pneumoniae* D39 (Δ*cps2A’*-Δ*cps2H’*) | [62] |
| D39 *hlpA::mKate2* | *mKate2*-expressing capsulated D39 (serotype 2) *S. pneumoniae,* Cam^R^ | [9] |
| D39 *∆cps hlpA::mKate2* | *mKate2*-expressing unencapsulated D39 *S. pneumoniae*, Cam^R^ | [9] |
| D39 *∆cps ∆ply hlpA::mKate2* | *mKate2*-expressing unencapsulated and pneumolysin-deficient D39 *S. pneumoniae*, Cam^R^, Kan^R^ | This study |
| **Oligonucleotide name** | **Sequence** | |
| AntibioticMarker_F | GAGGGAGGAAAGGCAGGA | |
| AntibioticMarker_R | CGCCGTATCTGTGCTCTC | |
| *ply*_5FLANK_F | CGATAAGGAAAAGATGAGCGCG | |
| *ply*_5FLANK_R | **TCCTGCCTTTCCTCCCTC**CTCCCTGATGGGTCAAGAGTTTC | |
| *ply*_3FLANK_F | **GAGAGCACAGATACGGCG**CTCTCTATCCTCAGGTAGAGGATAAGG | |
| *ply*_3FLANK_R | CTTCTTGTGCTTCTACGTCATCCAC | |
| *ply*_seq_F | CTCAATCCAGTTACCTGTCGCC | |

Cam = chloramphenicol, Kan = kanamycin resistance cassettes. For oligonucleotide sequences overlapping sequences used for isothermal assembly when generating gene knockout or genome integration constructs are shown in **bold**.

| **Gene** | **Identifier**  **(Ensembl)** | **Target location** | **Sequence (5’-3’)** | **Primers used for validation** | **Reference** |
| --- | --- | --- | --- | --- | --- |
| *atg5* | ENSDARG00000023396 | Exon 4 | GAGGTCGAACAACACACCAA(TGG) | F: AGATCTCTGCTTGCTTGGTT R: ACGAGGCAATCTACACACCT | [65] |
| *atg16l1* | ENSDARG00000099430 | Exon 2 | TTTGTGGAAGCGTCACGTTG(TGG) | F: TTGTGTCTGTCTGTTTGTGTGC  R: GGTGTATGATCTCCTCAAAGGT | [25] |
| *rubcn* | ENSDARG00000078752 | Exon 2 | CCCAACGTGTGGTCCCGGTA(TGG) | F: GAGCTCTGGAAGCTTCTGTATAA  R: CTGTTCGTGTTTGAGTCCGT | This study |
| *rubcn* | ENSDARG00000078752 | Exon 4 | TATAACCCTCGCCGAAGCTC(TGG) | F: GCTGATTTTTGCTGTGTCTCTG  R: CGTGTGGTGATACCGTTGTAAT | This study |
| *rubcn* | ENSDARG00000078752 | Exon 5 | TCGATGAGAGCCAGCATTCG(AGG) | F: GTGTCTGCAATGTTCCAGTGTC  R: TTTTTAAGAGGAGCGCATTAGC | This study |
| *tecpr1a* | ENSDARG00000062515 | Exon 2 | GTGTCCTCTTGGTAGCGGAT(GGG) | F: TGTACGGGAGGGTTTACAGTCT  R: CGGTTCATGTCTGTTCCTGTTA | This study |
| *tecpr1a* | ENSDARG00000062515 | Exon 5 | AGATCCCCTCAGATCAGACC(GGG) | F: TGCTCATGAAAACATTGTGTCTT  R: TTGAGTTCTCACCTTGCCCT | This study |
| *tecpr1b* | ENSDARG00000074086 | Exon 2 | GTCTCCTCCTGGTGTCGAAT(AGG) | F:  CTGGATCCGATCTGCTGACC  R:  CACCTCTGCACAACACACAG | This study |
| *tecpr1b* | ENSDARG00000074086 | Exon 4 | TCAGGGCTGGGAGTACGCTG(TGG) | F: CCAGTTATCATCCCATGCCTAT  R: TTATTGTCAACACTCCACAGCC | This study |

**Table S3.** CRISPR/Cas9 target sequences and primers for verification used in this study.

PAM site is indicated in brackets.

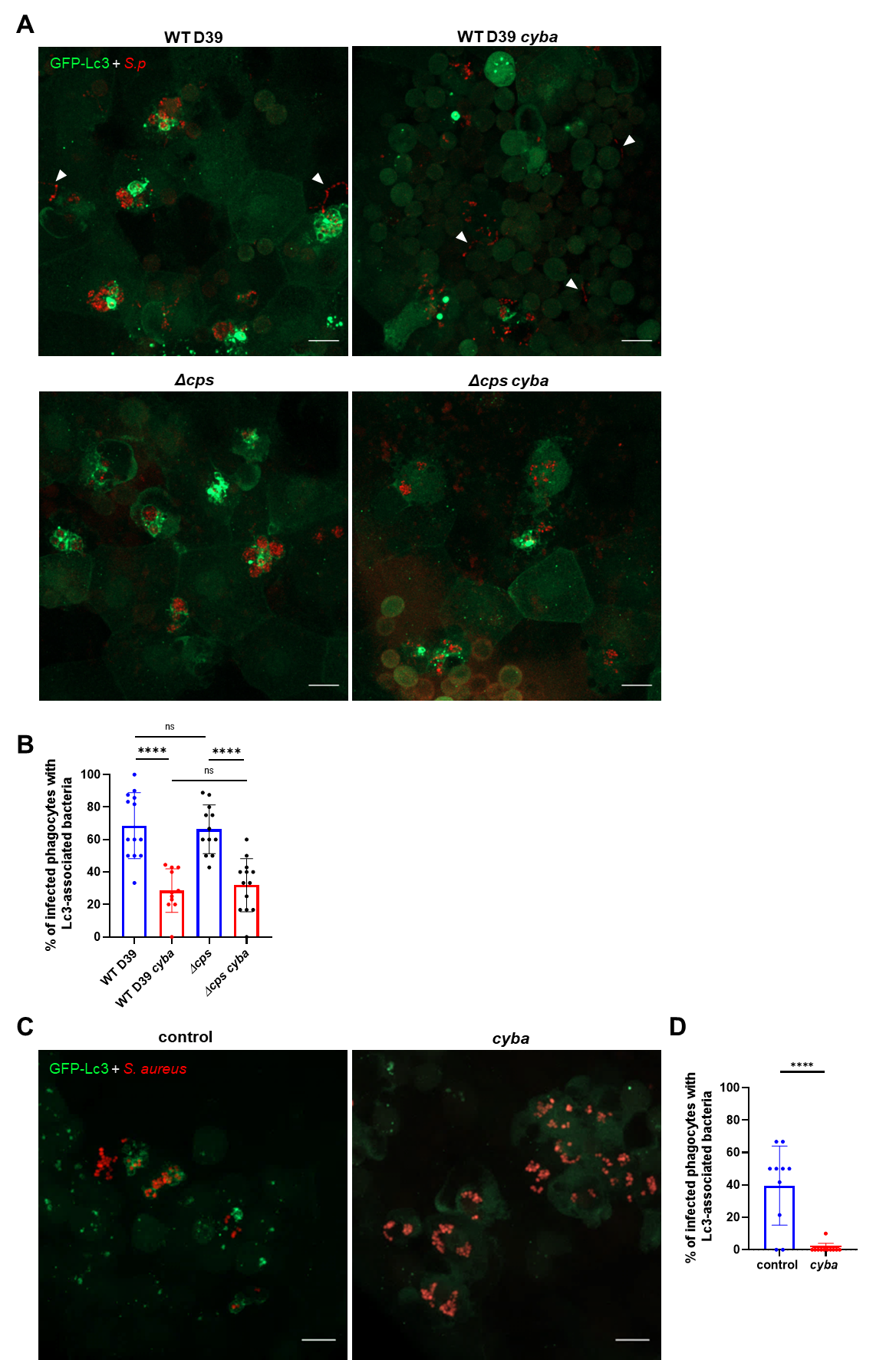

**Figure S1.** The autophagic response is triggered to intracellular encapsulated *S. pneumoniae*. (**A-B**) The autophagic response is triggered by intracellular encapsulated *S. pneumoniae*. (**A**) Representative confocal images of control or *cyba* knockdown *CMV:GFP-Lc3* larvae infected systemically with approximately 1600 CFU of mKate2-labeled encapsulated WT D39 (top panels) or unencapsulated D39 *Δcps* (bottom panels) *S. pneumoniae* and fixed at 2 hpi Scale bar: 10 µm. Arrowheads indicate non-phagocytosed bacteria. (**B**) Quantification of Lc3 associations with intracellular WT or *Δcps* D39 within infected phagocytes of fixed *CMV:GFP-Lc3* larvae. Data are shown as individual values ± standard deviation (SD). N≥11 zebraﬁsh larvae were analyzed. (**C-D**) NADPH oxidase is required for the Lc3-mediated response to *S. aureus* infection. (**C**) representative confocal images of control (left panel) or *cyba* knockdown (right panel) *CMV:GFP-Lc3* larvae infected systemically with approximately 1500 CFU of mCherry-labelled SH1000 *S. aureus* fixed 2 hpi. Scale bars: 10 µm. (**D**) Quantification of Lc3 associations with intracellular *S. aureus* within infected phagocytes of fixed *CMV:GFP-Lc3* larvae at 2 hpi. Data are shown as individual values ± standard deviation (SD). n≥10 zebraﬁsh larvae were analyzed.

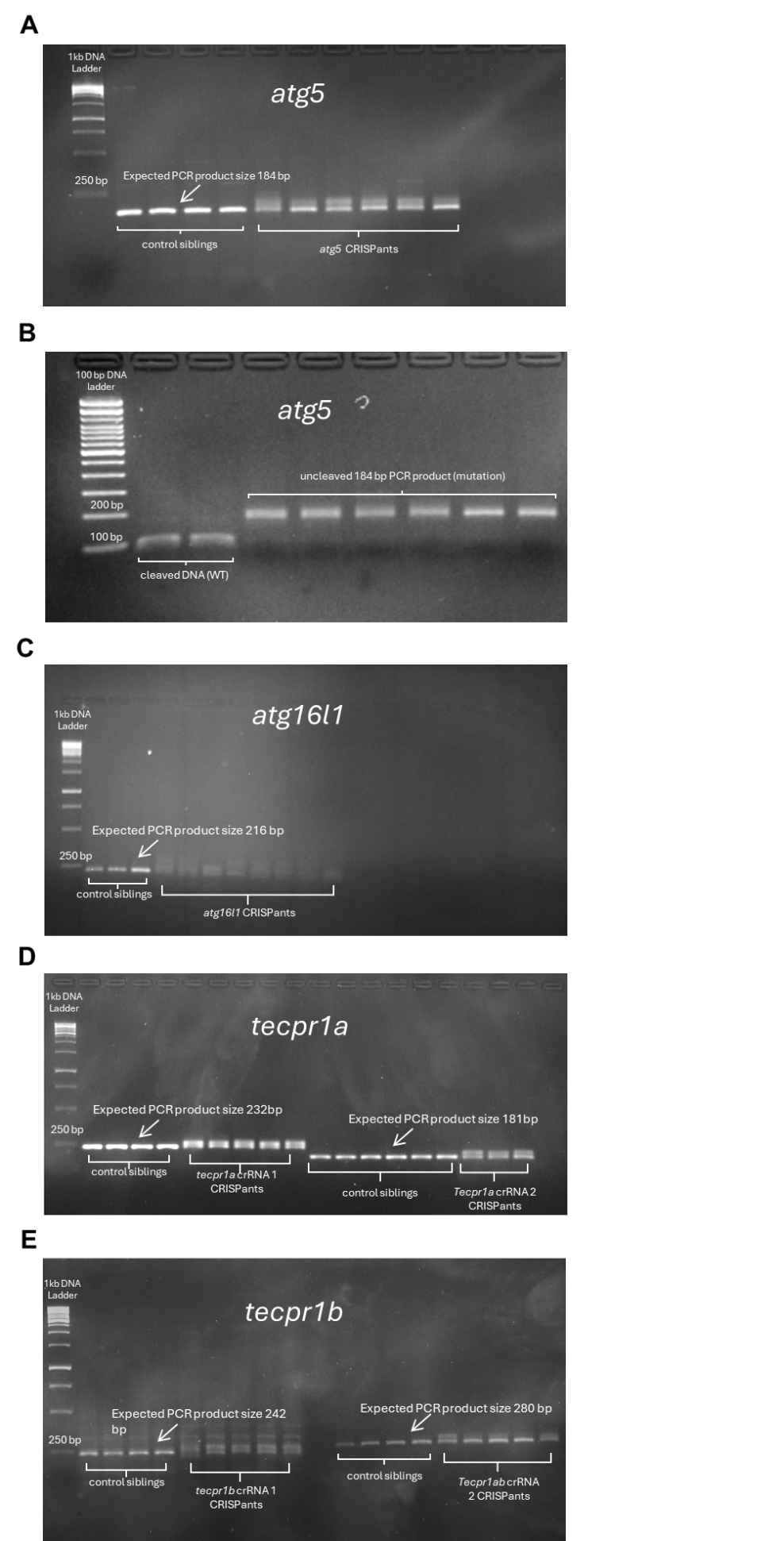

**Figure S2.** Efficacy verification of CRISPR-mediated knockdowns performed in this study. (**A, C, D, E**) Electrophoresis gel scans of polymerase chain reaction (PCR) products of control and (A) *atg5*, (C) *atg16l1*, (D) *tecpr1a* or (E) *tecpr1b* CRISPant zebrafish at 68 hpf used to determine the efficacy of mutagenesis. The expected PCR product sizes in unaffected larvae are labelled on scans. All tested CRISPant larvae showed mutagenesis as depicted by a smeared PCR product. (**B**) Electrophoresis gel scan of polymerase chain reaction (PCR) products of control and *atg5* CRISPants larvae digested with MslI restriction enzyme.

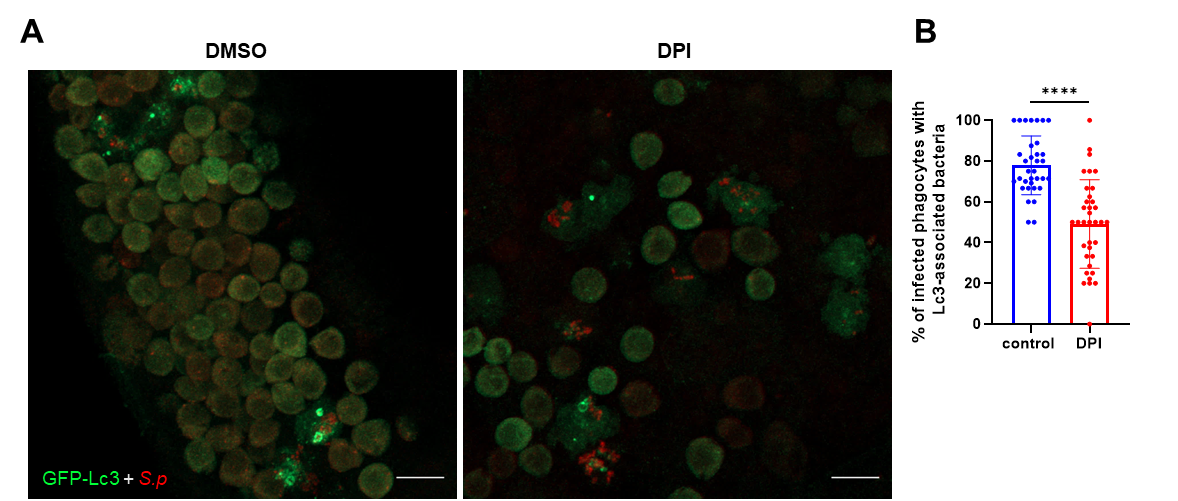

**Figure S3.** Chemical ROS inhibition results in reduction of autophagic response to pneumococci. (**A**) representative confocal images of DMSO (right panel) and DPI treated (right panel) *CMV:GFP-Lc3* larvae infected systemically with approximately 1600 CFU of mKate2-labeled D39 *Δcps* *S. pneumoniae* and fixed at 2 hpi. Scale bars: 10 µm. **B**) Quantification of Lc3 associations with intracellular *S. pneumoniae* within infected phagocytes of DMSO- and DPI- treated *CMV:GFP-Lc3* larvae fixed at 2 hpi. Data are shown as individual values ± standard deviation (SD). Data obtained from two independent experiments.

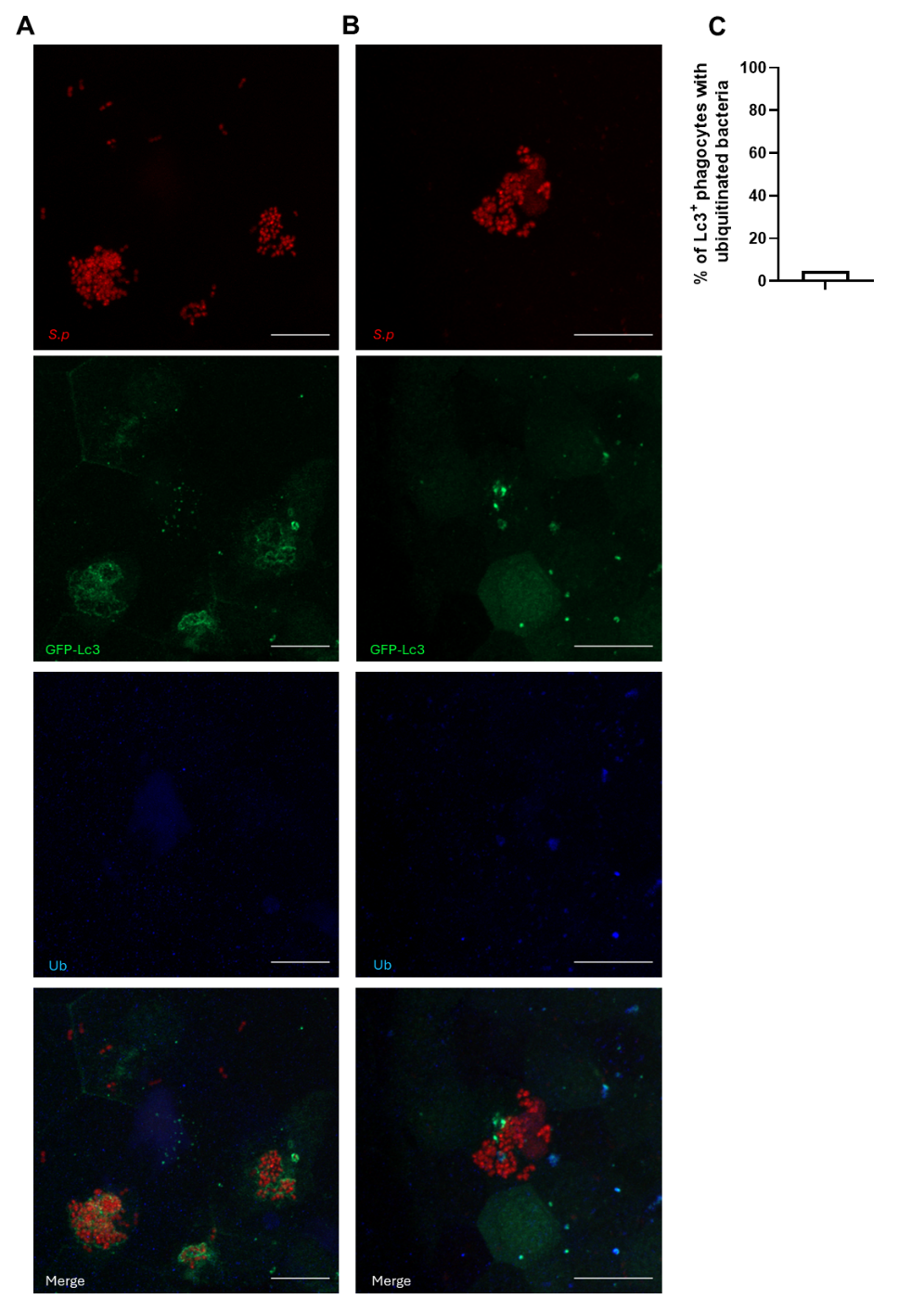

**Figure S4.** Ubiquitination is barely involved in autophagy response to intracellular *S. pneumoniae*. (**A-B**) Representative confocal images of (A) non-ubiquitinated and (B) ubiquitinated *S.p* clusters in GFP-Lc3 positive phagocytes at 2 hpi. Scale bars, 10 μm. (B) Quantification of the percentage of *S.p* clusters colocalized with GFP-Lc3 and ubiquitin at 2 hpi. Data obtained from two independent experiments.

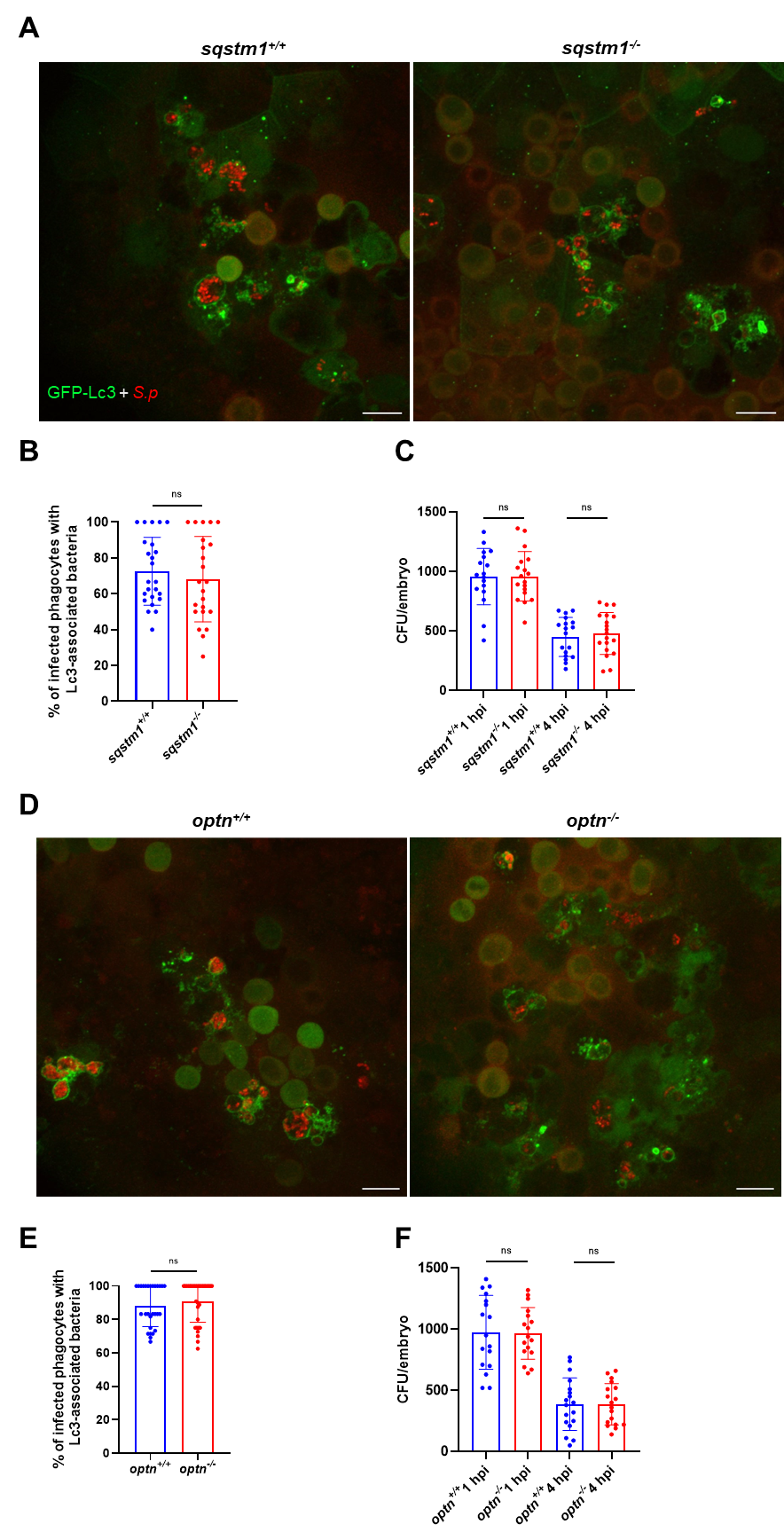

**Figure S5.** Single *sqstm1* and *optn* mutants are not deficient in Lc3-mediated response to pneumococci. **A, D**) representative confocal images of indicated mutant (right panel) or their respective wild-type siblings (left panel) of *CMV:GFP-Lc3* larvae infected systemically with approximately 1600 CFU of mKate2-labeled D39 *Δcps* *S. pneumoniae* and fixed at 2 hpi Scale bars: 10 µm. **B, E**) Quantification of Lc3 associations with intracellular *S. pneumoniae* within infected phagocytes of fixed *CMV:GFP-Lc3* wild type and indicated mutant larvae. Data are shown as mean ± standard deviation (SD). Data obtained from two independent experiments. **C, F**) Numbers of viable colony forming units (CFUs) over time of indicated mutant and their respective wild-type siblings upon infection with approximately 1600 CFU of *S. pneumoniae*. Data obtained from two independent experiments.

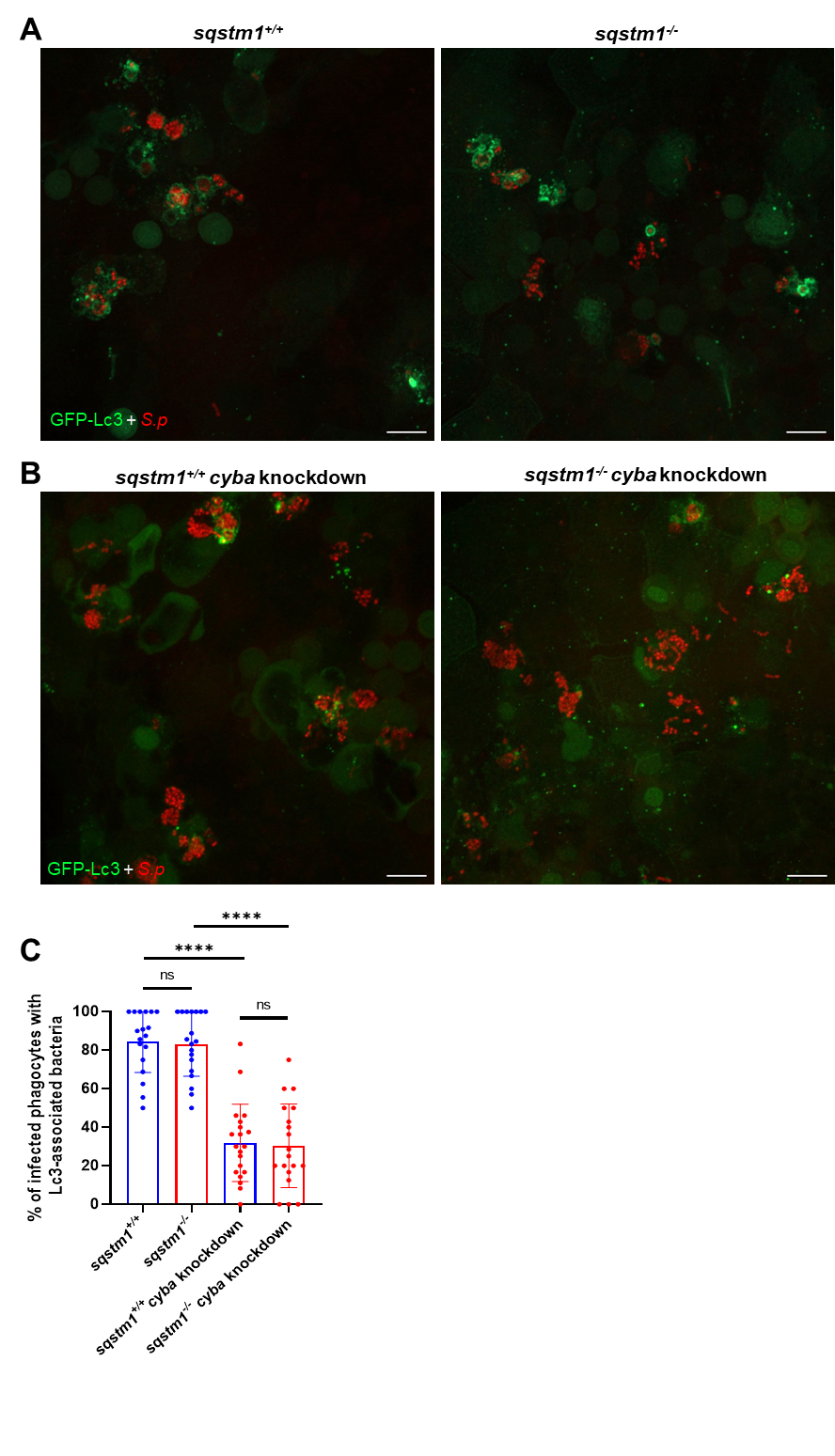

**Figure S6.** Knockdown of *cyba* does not result in an additional reduction of the autophagic response in *sqstm1 ^-/-^* mutant larvae. **A**) representative confocal images of *sqstm1*^+/+^ (left panel) and *sqstm1* ^-/-^mutant (right panel) *CMV:GFP-Lc3* larvae infected systemically with approximately 1600 CFU of mKate2-labeled D39 *Δcps* *S. pneumoniae* and fixed at 2 hpi. **B**) representative confocal images of *cyba* knockdown *sqstm1*^+/+^ (left panel) and *cyba* knockdown *sqstm1* ^-/-^mutant (right panel) *CMV:GFP-Lc3* larvae infected systemically with approximately 1600 CFU of mKate2-labeled D39 *Δcps* *S. pneumoniae* and fixed at 2 hpi Scale bars: 10 µm. **C**) Quantification of Lc3 associations with intracellular *S. pneumoniae* within infected phagocytes of fixed *sqstm1*^+/+^, *sqstm1*^+/+^  *cyba* knockdown, *sqstm1*^-/-^, and *sqstm1*^-/-^ *cyba* knockdown larvae in *CMV:GFP-Lc3* background. Data are shown as individual values ± standard deviation (SD). ≥18 larvae were analyzed.

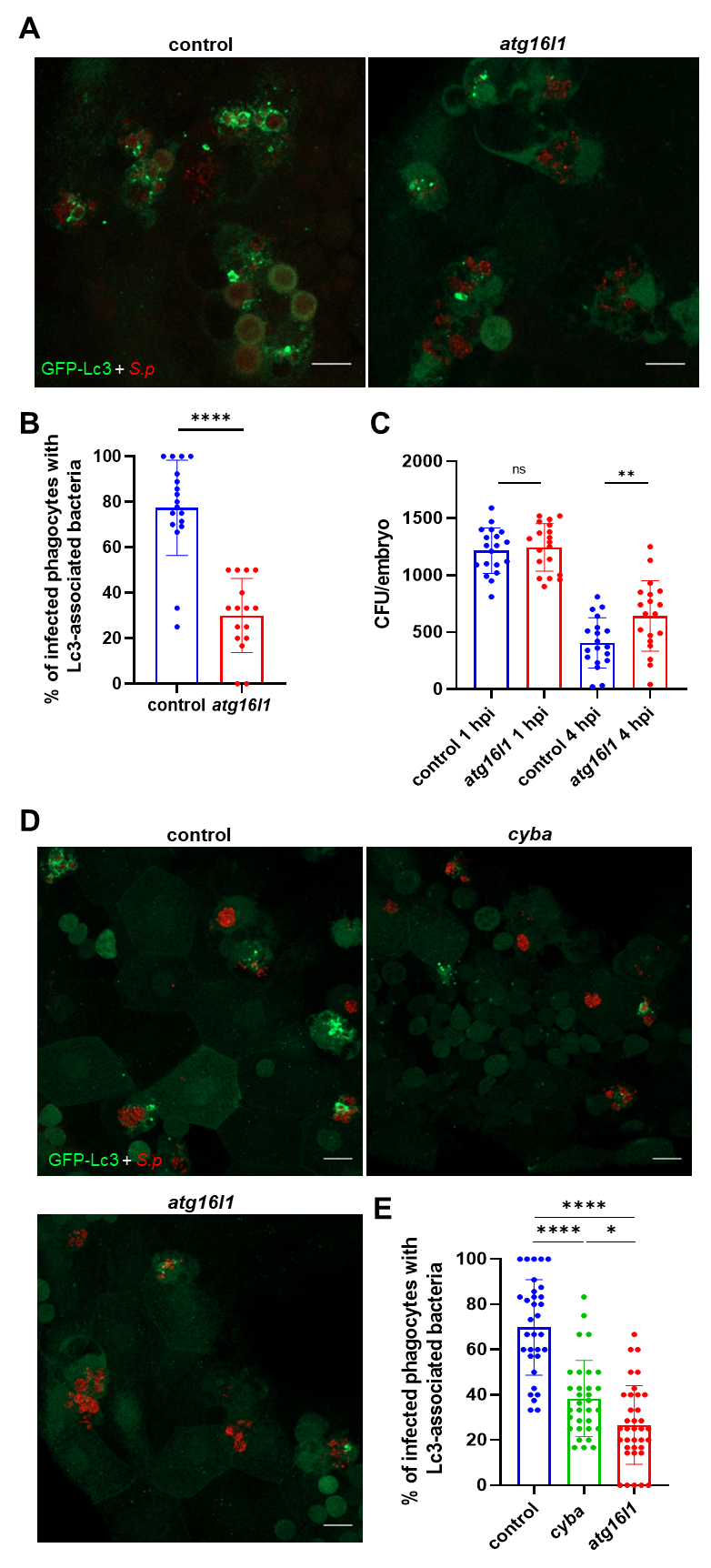

**Figure S7.** Knockdown of *atg16l1* results in partial reduction of the autophagic response to pneumococci larvae. (**A**) representative confocal images of control (left panel) and *atg16l1* knockdown (right panel) *CMV:GFP-Lc3* larvae infected systemically with approximately 1600 CFU of mKate2-labeled D39 *Δcps* *S. pneumoniae* and fixed at 2 hpi. Scale bars: 10 µm. (**B**) Quantification of Lc3 associations with intracellular *S. pneumoniae* within infected phagocytes of fixed control and *atg16l1* knockdown larvae in *CMV:GFP-Lc3* background. Data are shown as individual values ± standard deviation (SD). ≥16 larvae were analyzed. (**C**) Numbers of viable colony forming units (CFUs) over time of control and *atg16l1* knockdown larvae. Data obtained from two independent experiments. (**D**) representative confocal images of control, *cyba* or *atg16l1* knockdown *CMV:GFP-Lc3* larvae infected systemically with approximately 1600 CFU of mKate2-labeled D39 *Δcps* *S. pneumoniae* and fixed at 2 hpi. Scale bars: 10 µm. (**E**) Quantification of Lc3 associations with intracellular *S. pneumoniae* within infected phagocytes of fixed control, *cyba* or *atg16l1* knockdown larvae. Data are shown as individual values ± standard deviation (SD). Data obtained from two independent experiments.

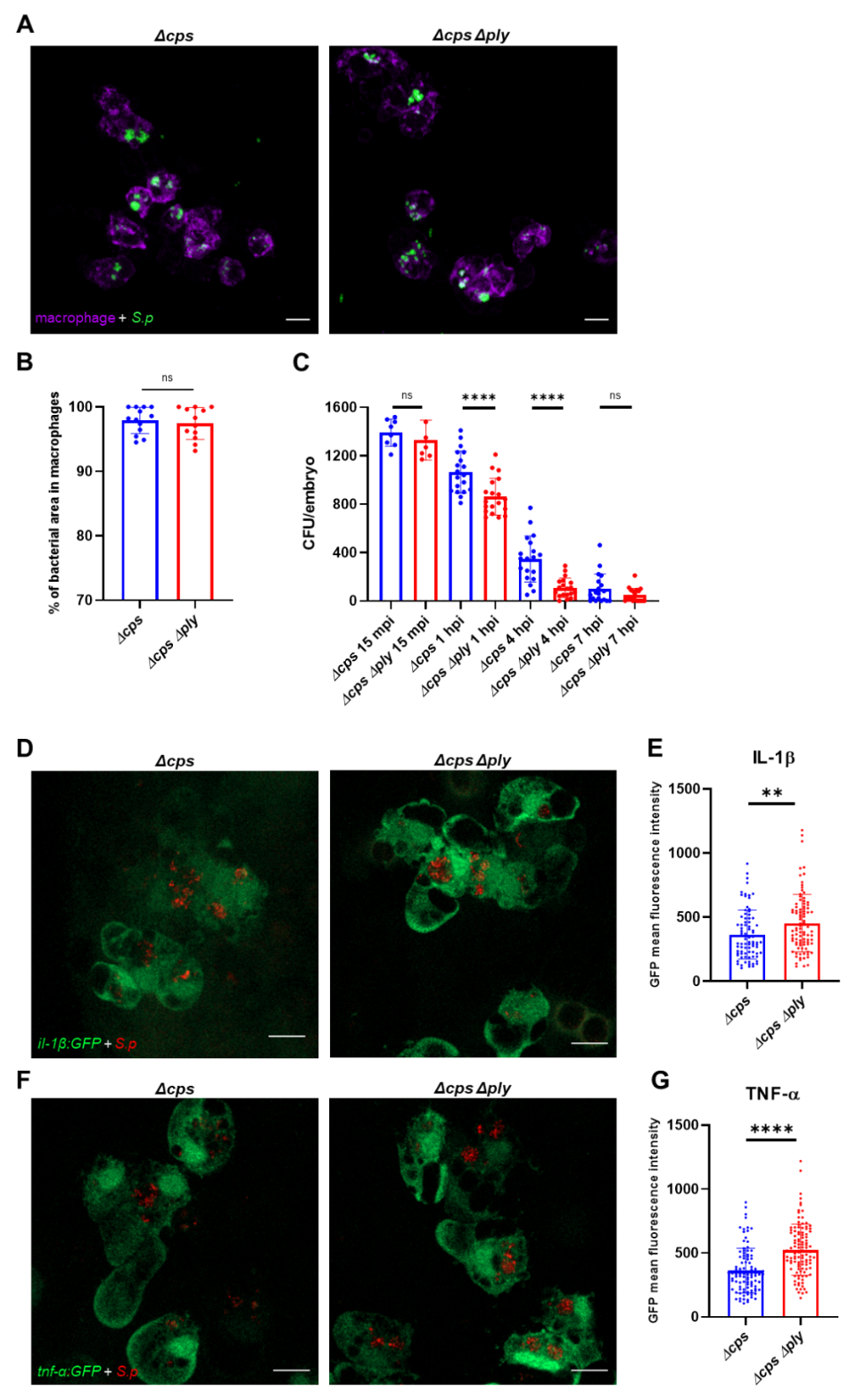

**Figure S8.** Lack of pneumolysin results in stronger proinflammatory response to *S. pneumoniae* and more effective elimination of pneumococci. (**A**) representative confocal images of *mpeg:mCherry* larvae infected systemically with approximately 1600 CFU of Alexa fluor 647 prestained, mKate2-labeled PLY-positive D39 *Δcps* (left panel) or PLY-negative D39 *Δcps* *Δply* (right panel) *S. pneumoniae* and fixed at 2 hpi. Scale bars: 10 µm. (**B**) Quantification of bacterial are within mCherry-labeled macrophages. Scale bars: 10 µm. n≥12 larvae were analyzed. (**C**) Numbers of viable colony forming units (CFUs) over time of *mpeg:mCherry* larvae upon infection with approximately 1600 CFU PLY-positive and PLY-negative *S. pneumoniae*. Data obtained from two independent experiments. (**D, F**) representative confocal images of *il1b:GFP* (D) or *tnfa:GFP* (F) larvae infected systemically with approximately 1600 CFU of mKate2-labeled PLY-positive D39 *Δcps* (left panels) or PLY-negative D39 *Δcps* *Δply* (right panels) *S. pneumoniae* and fixed at 2 hpi. Scale bars: 10 µm. (**E, G**) Quantification of GFP mean fluorescence signal within individual infected phagocytes of fixed *il1b:GFP* (E) or *tnfa:GFP* (G) larvae. Data are shown as individual values ± standard deviation (SD). Data obtained from two independent experiments.

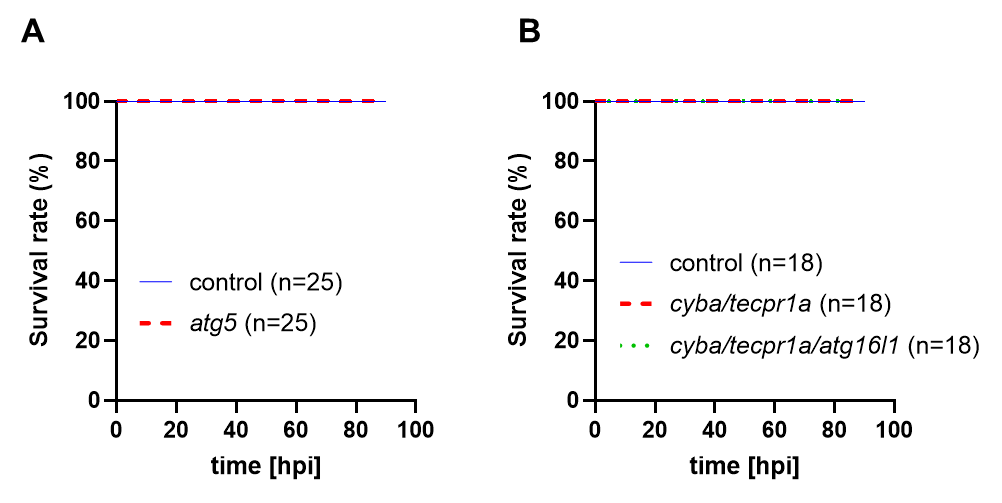

**Figure S9.** Lack of autophagy does not affect the survival of pneumococcus-infected larvae. (**A**) survival of control and *atg5* knockdown larvae following infection at 36 hpf with approximately 1600 CFU of mKate2-labeled D39 *Δcps* *S. pneumoniae.* (**B**) survival of control, *cyba/tecpr1a* or *cyba/tecpr1a/atg16l1* knockdown larvae following infection at 36 hpf with approximately 1600 CFU of mKate2-labeled D39 *Δcps* *S. pneumoniae*.
